# Effect of Alcohol and Cocaine Abuse on Neuronal and Non-Neuronal Cell Turnover in the Adult Human Hippocampus

**DOI:** 10.1101/2024.09.10.612190

**Authors:** Kanar Alkass, Embla Steiner, Samuel Bernard, Tara Wardi Le Maître, Gopalakrishnan Dhanabalan, Mikael Landén, Kirsty Spalding, Jonas Frisén, Deborah C Mash, Henrik Druid

**Author notes:** These authors contributed equally to the study (Kanar Alkass, Embla Steiner).

## Abstract

Clinical studies on humans with a history of chronic abuse of alcohol or cocaine show cognitive impairments associated with hippocampal atrophy. Adult hippocampal neurogenesis is a process important for memory formation and has been shown to be impaired by alcohol and cocaine in rodent models. It has thus been suggested that a reduction in adult neurogenesis may contribute to cognitive dysfunctions seen in patients with abuse. In addition, reduced adult neurogenesis has been suggested to play a role in the pathology of addiction vulnerability. We have previously demonstrated persistent adult hippocampal neurogenesis throughout life by measuring ^14^C concentrations in genomic DNA, incorporated during cell division, in a mixed cohort of subjects. In this study, we use the same strategy to assess the extent of cell turnover of neuronal and non-neuronal cells in the hippocampus of humans with known history of alcohol and cocaine abuse and compare these with healthy controls. We find that there is significant neuronal and non-neuronal turnover in healthy controls, as well as in individuals with long term alcohol use or cocaine use. Using mathematical modelling, we compare the extent of cell turnover of neurons and non-neuronal cells and did not find any significant difference between healthy controls and the two addiction groups. While we cannot exclude scenarios of altered adult neurogenesis over shorter periods of time, our data does not support the theory of low neurogenesis as a mechanism of addiction vulnerability.

## Introduction

Substance abuse is associated with several health risks and social problems. Both cocaine and alcohol intake results in damage of the structure and function of brain areas associated with cognition and motor skills^1,2^. Findings from neuroimaging, physiological and neuropsychological studies of alcoholics illustrate that the frontal cortex, the limbic system, and cerebellum are particularly vulnerable to damage^3,4^. Hippocampus, being part of the limbic system, is a brain region involved in declarative memory, higher cognitive functions, and regulation of emotional behaviors. Hippocampus is also a part of the memory/conditioning neuronal circuitry which is considered to be important for drug-induced reinforcement in animals as well as for cravings and relapses in humans^5,6^.

A special feature of the hippocampus is that new neurons, generated from multi-potent progenitor cells, integrate throughout life in the granular cell layer of the dentate gyrus (DG) in a process called adult neurogenesis^7^. This process represents an important form of hippocampal plasticity that is critical for declarative memory consolidation in experimental animals^8,9^. Dysfunction in adult neurogenesis has been suggested to contribute to diseases such as Alzheimer’s disease, major depression, and schizophrenia as well as drug addiction^10–15^. Learning and conditioning contributes to drug tolerance, dependence and relapse^16^ and substance abuse is frequently associated with cognitive disorders, indicative of hippocampal dysfunction.

Adult hippocampal neurogenesis and its role in drug addiction has been studied in the context of the rodent brain. Alcohol targets multiple steps of adult neurogenesis, including cell proliferation, neuronal differentiation, cell migration, cell survival and circuitry integration^17–19^. Cocaine administration reduces cell proliferation in the adult rat hippocampus^20,21^ without altering the survival or dendritic maturation of newly generated cells^22^. Furthermore, withdrawal from cocaine or alcohol normalizes proliferation deficits and enhances the maturity of adult generated neurons^23,24^.

Few studies have investigated how human adult hippocampal neurogenesis is affected by alcohol and cocaine, mainly due to the lack of methods to study this process in the human brain. MRI studies have shown structural changes in the hippocampus in individuals with alcohol or cocaine use^25–28^, and studies using stereology techniques have demonstrated neuronal loss in the hippocampus^29,30^. However, these findings cannot be directly correlated to alterations in adult neurogenesis. Analysis of molecular markers of proliferation and neural precursor cells in donors with on-going alcohol abuse show a reduction of neural stem/progenitor cells and immature neurons in the DG^31^. This provides a good view of the number of cells in the cycle at a given time, but it does not provide the number of mature cells generated or added to an existing network over lifetime.

Retrospective birth dating of cells by measuring ^14^C concentrations in genomic DNA provides a unique possibility to study neuronal generation in the hippocampus throughout life^32^. Utilizing the dramatic increase of atmospheric ^14^C caused by nuclear bomb tests during the cold war, we can measure the average concentration of ^14^C in the genomic DNA incorporated during cell division. Thereby, the ^14^C concentration acts as a natural cell division marker. By measuring the average ^14^C in millions of cells this concentration reflects the average cell age at the date of collection. Mathematical modelling allows us to test different biological scenarios of cell turnover, the process of dying cells being replaced by new cells, over a subject’s life and exclude models that are not compatible with the measured ^14^C data. Previously, this strategy has been used to study adult hippocampal neurogenesis in a mixed cohort, showing substantial neurogenesis throughout life in the human hippocampus^32^. Here, we report cell turnover estimations in the hippocampus in a carefully selected healthy control group and investigate whether significant changes occur in the dynamics of adult hippocampal neurogenesis throughout life in individuals with alcohol or cocaine abuse.

## Materials and Methods

### Tissue Collection

Hippocampus brain samples, including both the DG and the cornu ammonis areas 1 to 4, from chronic alcohol abusers and control subjects were collected by KI Donatum for the project from deceased donors subjected to forensic autopsy at the Department of Forensic Medicine in Stockholm. Consent was either obtained from the Swedish donation registry or from next-of-kin. In both cases, the relatives are contacted by the forensic investigator at the Department of Forensic Medicine. Ethical permission for this study was granted by Regional Ethics Committee of Sweden (EPN Dnr 2010/313-31/3 and 2018/689-32). Samples from deceased alcohol addicts and cocaine addicts, and from control subjects were also obtained from the Miami Brain Endowment Bank after obtained consent. Ethical permission was approved by the institutional review board of the University of Miami Miller School of Medicine (19920580 (CR00006788)). Whole hippocampi were dissected and analyzed. Brain tissue was frozen and stored at −80°C until further analysis. Characterization of subjects regarding alcohol consumption, legal and illegal drug use, medical condition, physical activity, and more, was performed by structured interviews with relatives, and by perusing the forensic pathology investigations, police reports, and medical records, when available as well as from the Swedish national patient register and prescribed drug register. The result of 70 of the 144 measured ^14^C samples have previously been reported^32^, but are now subject to extended mathematical modeling.

### Nuclei Isolation

Tissue samples of 2-6 grams were thawed and homogenized using a Dounce Homogenizer. About 1 gram of tissue was homogenized in 10 ml lysis buffer (0.32 M sucrose, 5 mM CaCl_2,_ 3 mM magnesium acetate, 0.1 mM EDTA, 10 mM Tris-HCl [pH 8.0], 0.1% Triton X-100, and 1 mM DTT). Each 1 gram of homogenized samples was suspended in 20 ml of sucrose solution (1.7 M sucrose, 3 mM magnesium acetate, 1 mM DTT, and 10 mM Tris-HCl [pH 8.0]), layered onto a cushion of 10 ml sucrose solution, and centrifuged at 13000 rpm for 2 hours and 20 minutes at 4°C. The isolated nuclei were resuspended in nuclei storage buffer (10 mM Tris [pH = 7.2], 2 mM MgCl_2_, 70 mM KCl, and 15% sucrose) for consecutive immunostaining and flow cytometry analysis.

### Immunostaining and FACS Sorting and Analysis

Isolated nuclei were stained with mouse NeuN (A-60) (Millipore, 1:1000), which was directly conjugated to Alexa 647 using the Alexa Flour 647 Antibody Labeling Kit (Invitrogen). Fluorescence-activated cell sorting (FACS) was performed with a Vantage (Becton Dickinson) or a MoFlo XDP (Beckman Coulter) and influx (BD Bioscience) flow cytometer. The FACS gating strategy for sorts is shown in Supplementary Fig. 1. Nuclei pellets were collected after centrifugation and stored at -80°C until DNA extraction was performed. The FACS purity was later used to purity-correct measured ^14^C values as previously described (see Supplementary material)^32^.

### DNA Purification

All experiments were carried out in a clean room (ISO8) to prevent any carbon contamination of the samples. All glassware was prebaked at 450°C for 4 hr. DNA isolation was performed according to a modified protocol from Miller et al., 1988. Then, 500 μl DNA lysis buffer (100 mM Tris [pH 8.0], 200 mM NaCl, 1% SDS, and 5 mM EDTA) and 6 μl Proteinase K (20 mg/ml) were added to the collected nuclei and incubated overnight at 65°C. RNase cocktail (Ambion) was added and incubated at 65°C for 1 hour. Half of the existing volume of 5 M NaCl solution was added, and the mixture was agitated for 15 seconds. The solution was spun down at 13.000 rpm for 3 minutes. The supernatant containing the DNA was transferred to a 12 ml glass vial. Three times of the volume of absolute ethanol were added, and vial was inverted several times to precipitate the DNA. The DNA precipitate was washed three times in DNA-washing solution (70% ethanol [v/v] and 0.1 M NaCl) and transferred to 500 μl DNase-and RNase-free water (Gibco and Invitrogen, respectively). The DNA was quantified, and DNA purity was verified by UV spectroscopy (NanoDrop).

### Accelerator Mass Spectrometry

All Accelerator mass spectrometry analyses were performed without knowledge of the sample’s age and origin to prevent bias. Purified DNA samples suspended in water were lyophilized to dryness. To convert the DNA sample into graphite, excess CuO was added to each dried sample, and the tubes were evacuated and sealed with a high-temperature torch. Tubes were placed in a furnace set at 900°C for 3.5 hours to combust all carbon to CO_2_. The evolved CO_2_ was purified, trapped, and reduced to graphite in individual reactors at 550°C for 6 hours, using iron as catalyst. Graphite targets were either measured at the Center for Accelerator Mass Spectrometry at the Lawrence Livermore National Laboratory or the Department of Physics and Astronomy, Ion Physics, of Uppsala University. Large CO_2_ samples (>100 μg) were split, and δ^13^C was measured by stable isotope ratio mass spectrometry, which resulted in a δ^13^C correction of –22.3‰ ± 0.5‰ (1 SD), which was applied for all samples. Corrections were made for background contamination introduced during sample preparation, as previously described^33,34^. All ^14^C data are reported as fraction modern.

### Statistics and Mathematical Modeling

The measured ^14^C value in our data is the average ^14^C concentration in the DNA from a population of cells. This value reflects the average age of the cell population, which depends on birth and death processes within that population over time. Mathematical modelling allows us to explore different biological scenarios and exclude those that are not compatible with our data. We further developed and extended previously described models based on birth and death processes (for detailed description see supplementary material)^32,35^. The mathematical models result in a cell age distribution matrix, which is then integrated over the atmospheric bomb curve to estimate a ^14^C value for that cell age distribution. To test if a scenario is compatible with our data, we performed *global modelling,* where estimated ^14^C values are fitted to measured ^14^C values. The sum of squared errors is estimated and determines the goodness of fit.

To determine the possible parameter values, we used Markov Chain Monte Carlo algorithms. This Bayesian strategy simulates thousands of parameter values for each model, testing the goodness of fit for each parameter values that are tested. We determined the best scenario by calculating the Akaike information criterion (AIC) for the best fit parameter values from each simulation. Likelihood ratio test was used to accept or reject a more complex parameter (see Table 2, 4 and 5 for model summaries, AIC, and best parameter estimates). If the parameter estimates from the best-fitted scenario resulted in estimates that made the complex model similar to a less complex model, the less complex model was chosen. If the values were biologically impossible, we could accept the model, but were careful to interpret the estimated values. We also performed *individual modelling* of each datapoint based on the best model chosen from global fits. This allowed us to estimate the optimal parameter value matching the measured ^14^C value, enabling direct comparison between the test groups.

## Results

### Measurement of ^14^C reveals significant turnover of hippocampal neurons and non-neuronal cells in healthy control, alcohol and cocaine groups

Cell turnover is the process of a cell dying and being replaced by a new cell generated through division of a progenitor cell. The retrospective birth dating method allows us to investigate turnover processes in cell populations, due to the dynamic atmospheric ^14^C concentrations that are incorporated in the DNA during cell division. The measured ^14^C from genomic DNA reflects the average concentration in the population of cells remaining at collection and can be translated to an average cell age. If there is no postnatal cell turnover, the cells at collection will only have incorporated atmospheric concentrations around the time of birth (Fig. 1a). If there is significant cell turnover, different ^14^C concentrations will be incorporated throughout life, resulting in a measured ^14^C concentration either higher or lower than the atmospheric concentration at time of birth (Fig. 1b).

**Figure 1:**
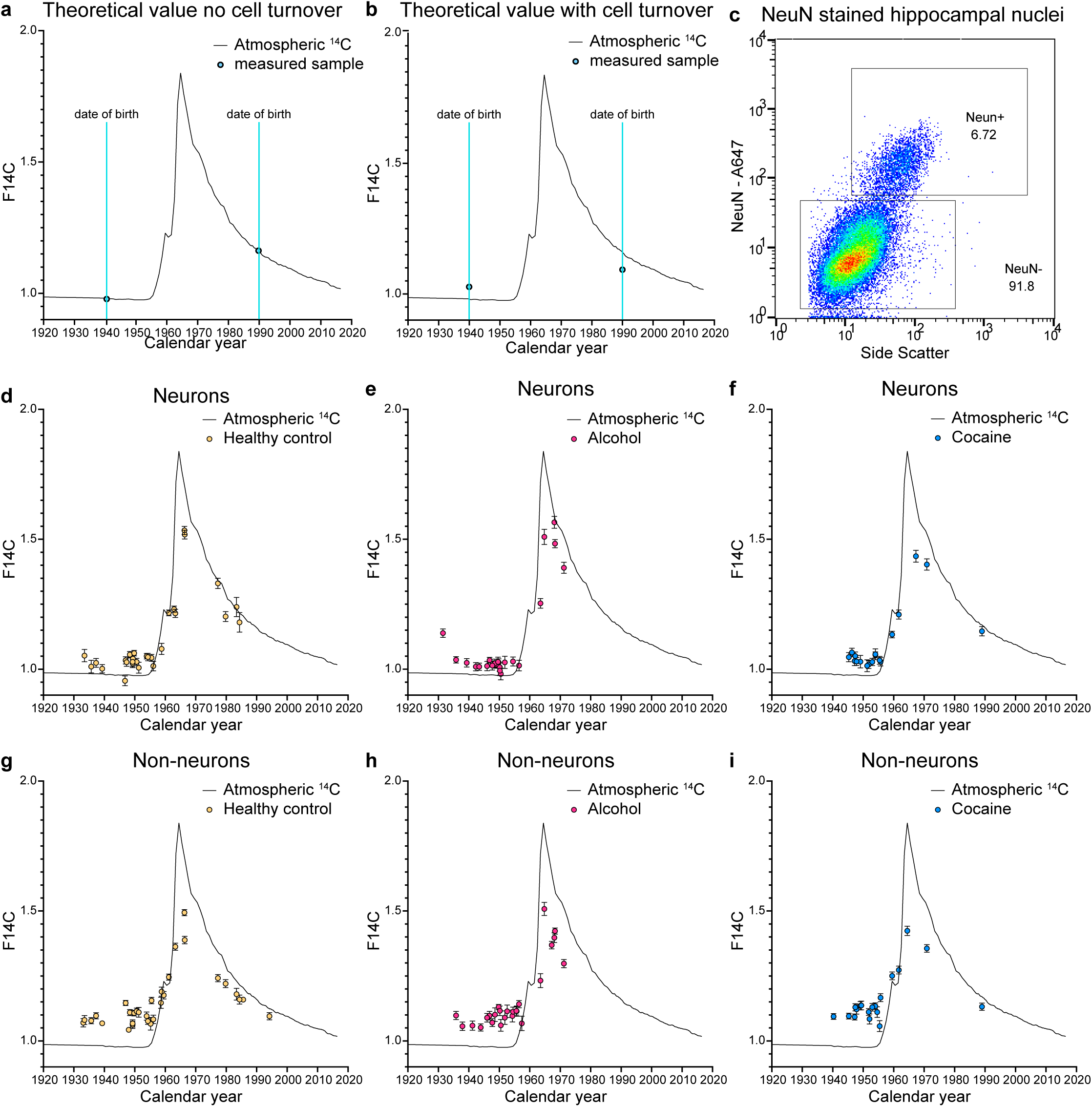
Measurement of ^14^C concentrations in genomic DNA. Theoretical measured ^14^C values if **(a)** there is not postnatal cell turnover or **(b)** there is postnatal cell turnover. **(c)** Gating of NeuN-positive neuronal nuclei and NeuN-negative non-neurons during fluorescence-activated cell sorting. Measured ^14^C in genomic DNA from hippocampal neurons and non-neuronal cells are shown for **(d** and **g)** healthy control group, **(e** and **h)** alcohol group and **(f** and **i)** cocaine group. Measured F14C, plotted along the calendar year at the time of birth of the individual. Black line represents the measurements of atmospheric F14C.

Neurons and non-neuronal cells were collected from the hippocampus, including the DG and the cornu ammonis (CA) 1-4 structures (Fig. 1c, Supplementary Fig. 1). Measurement of ^14^C concentration in DNA isolated from these cell populations, revealed significant cell turnover within both populations in the healthy control, alcohol and cocaine abuse groups (Supplementary Table 1, Fig. 1d-i). Directly comparing the ^14^C concentrations to estimate differences in cell turnover between groups is not possible due to the dynamic change in ^14^C, requiring the cohorts to be perfectly matched in date of birth and age, which was not possible in this study (Supplementary Fig. 2). Therefore, to compare cell turnover, we further developed previously described mathematical models to test different scenarios of turnover (see methods and Supplementary material)^32,35^.

### Mathematical modelling

The mathematical models simulate cell birth and death processes that affect the average age of a cell population, focusing on two concepts: cell turnover and selective cell death. Cell turnover is the process of replacing an older cell with a new cell, while leaving the total cell number constant and tends to maintain a cell population young. Selective cell death involves a dying cell of a specific age without being replaced by a new cell, reducing the cell numbers and affecting the average age. Within the hippocampal neuronal population, the impact of cell turnover on the average cell age depends on 1. The proportion of CA to DG neurons, as adult neurogenesis is limited the DG but we collect NeuN-positive neuronal nuclei from both the CA and DG areas. 2. The number of new neurons exchanged per year and 3. The dynamics of adult neurogenesis over time in the healthy and pathological brain (Supplementary Fig. 3a). Studies indicate that adult neurogenesis declines with age and in pathological conditions, affecting the number of newborn cells^32,36,37^. Less is known about potential selective cell death within hippocampal neurons. Stereology data suggest a selective loss of CA neurons with age, leading to the disappearance of older neurons (Supplementary Fig. 4)^38–42^. There are also studies suggesting selective loss of CA neurons after alcohol abuse which could affect the average age^29^.

We developed model scenarios, to describe these different processes that affect the average age. The two main components we used in our scenarios were 1. The proportion of cells subject to turnover, where we have a parameter referred to as the *renewing fraction* and 2. The turnover dynamics, where we estimate the exchange within this *renewing fraction* with a parameter referred to as *turnover rate*. We then added complexities with additional parameters of cell turnover dynamics over time and selective cell loss (Supplementary Fig. 3b).

To test which modelled scenario best fit our data, we performed *global modelling*, simulating thousands of parameter values and assessing how well these models fit our measured ^14^C data using Markov Chain Monte Carlo algorithms. The models with best fit were selected using AIC. For comparing the cell turnover between the different groups, we also performed *individual modelling* where we estimated the individual *turnover rates* for each sample. From these individual *turnover rates* we modeled an average cell age and compared the difference between individual person age and cell age, further referred to as *delta age*, across the three groups.

### Cell turnover in the non-neuronal population of alcohol and cocaine addicts could not be distinguished from the turnover in healthy controls

We modelled 8 biological scenarios for the non-neuronal population (Supplementary Table 2). Since the non-neuronal population represents a heterogenous group of cells and stereology data was not available, we did not model any scenarios of selective death or reduced *renewing fraction*. The best accepted model for all three groups was scenario 5, which estimates a *renewing fraction*, in which cells turn over with a constant *turnover rate* (Supplementary Table 2 and 4).

The global fit parameter estimates for this scenario were similar between the three groups. The initial *renewal fraction* was 0.65 for the control group, 0.60 for the alcohol group and 0.61 for the cocaine group. The annual *turnover rate* was 3.1% for the control group, 3.6% for the alcohol group and 4.3% for the cocaine group (Supplementary Fig. 5a-c). Thus, the global fit parameter estimation was similar between healthy controls, the alcohol group and the cocaine group.

Next, we modeled the individual annual *turnover rates* for all the samples by fixing the *renewal fraction* to the healthy control *global modelling* estimate of 0.65. There was a significant correlation between annual *turnover rates* and age for the healthy control (F[1, 25]= 22.49, R2= 0.5, p< 0.0001, linear regression) and cocaine (F[1, 14]= 5.78, R2= 0.3, p= 0.0307, linear regression) groups, but not the alcohol group (F[1, 13]= 0.78, R2= 0.06, p= 0.3939, linear regression). There was no significant difference between the individual annual *turnover rates* when comparing the three groups (Fig. 2a, F[5, 52]= 3.990 R2= 0.28, P= 0.0039, age was significant: F[1, 52]= 16.66, p= 0.0002. Group effect was not significant: F[2, 52]= 0.2688, p= 0.7654, multiple linear regression)

**Figure 2:**
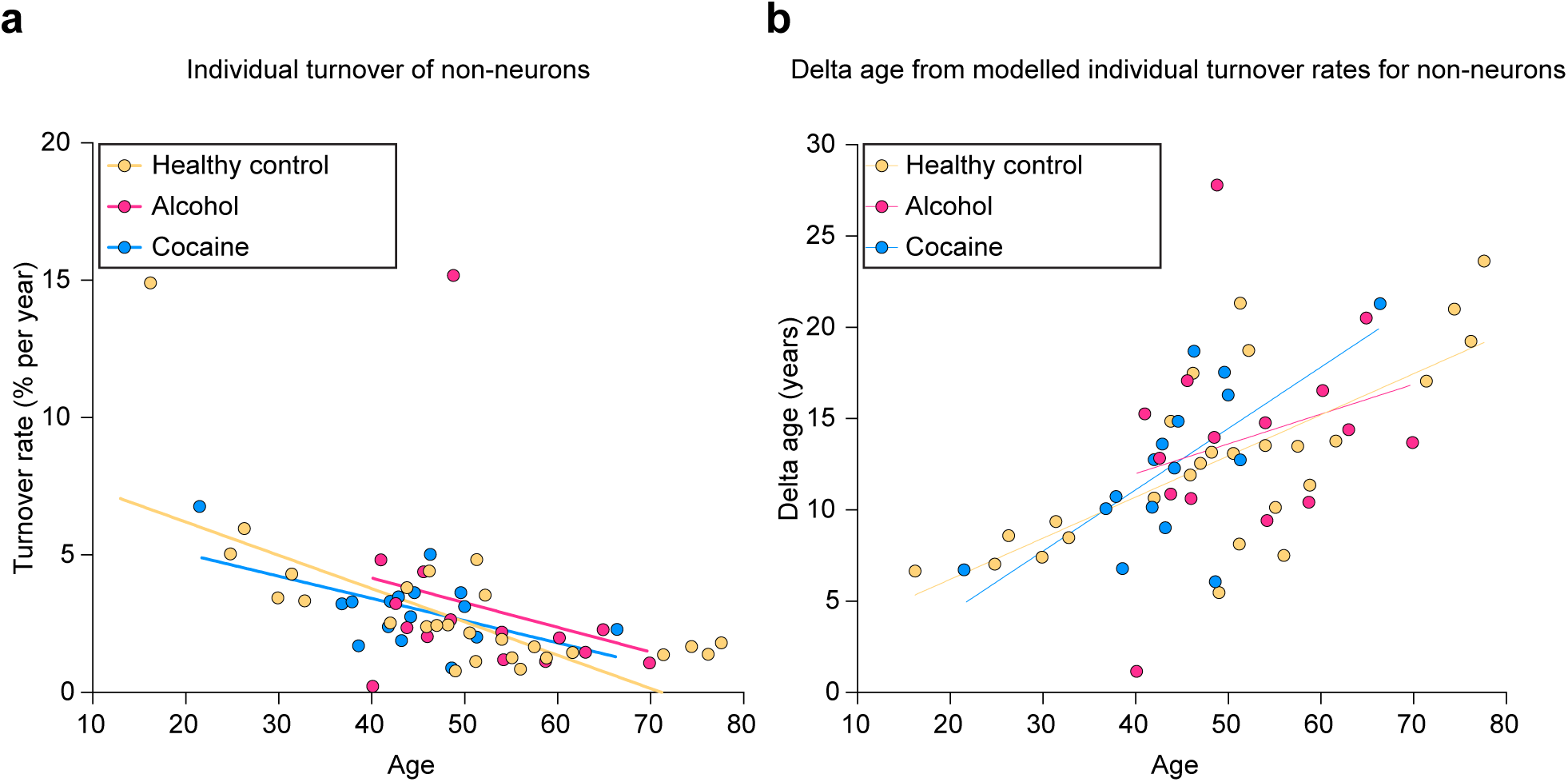
Individual modelling of non-neuronal cells. **(a)** Correlation between annual turnover rate of non-neurons and age. A significant correlation between annual turnover rates and age was found for the healthy control and cocaine groups, but not the alcohol group (Healthy control: F[1, 25]= 25.49, R2= 0.5, p< 0.0001, Alcohol: F[1, 13]= 0.78, R2= 0.06, P= 0.3939; Cocaine: [F(1, 14]= 5.78, R2= 0.3, P=0.0307, linear regression). There was no significant difference between the individual annual turnover rates when comparing the three groups (F[5, 52]= 3.990, R2= 0.28, p= 0.0039, age was significant: F[1, 52]=16.66, p = 0.0002. Group effect was not significant: F[2, 52]= 0.2688, p= 0.7654, multiple linear regression) **(b)** Correlation between non-neuronal delta age and age. There was a significant correlation between delta age and age for the healthy control and cocaine group but not the alcohol group (Healthy control: F[1, 25]= 25.00, R2= 0.5, p<0.0001; Alcohol: F[1, 13]= 0.98, R2= 0.07, p=0.34, Cocaine: F[1, 14]= 13.39, R2= 0.49, p= 0.0026, linear regression). There was no significant difference between the three groups (F[3, 55]= 201.7, R2= 0.92, p< 0.0001. Age was significant: F[1, 55]= 278.0, p< 0.0001. Group effect was not significant: F[2, 55]= 0.5329, p= 0.5899, multiple linear regression). Delta age= the difference in the person’s chronological age and the average cell age in years.

Using the individual *turnover rates* we modelled the average cell age and calculated the *delta age*. There was a significant correlation between non-neuronal *delta age* and age for the healthy control (F[1, 25] = 25.00, R2=0.5, P<0.0001, linear regression) and cocaine (F[1, 14]= 13.39, R2= 0.49, p= 0.0026, linear regression) groups, but not the alcohol group (F[1, 13]= 0.98, R2= 0.07, p=0.34, linear regression). There was no significant difference between the three groups (Fig. 2b, F[3, 55]= 201.7, R2= 0.92, p < 0.0001. Age was significant: F[1, 55]= 278.0, p< 0.0001. Group effect was not significant: F[2, 55]= 0.5329, p= 0.5899, multiple linear regression).

As we could not detect any significant difference in non-neuronal cell turnover between the three groups comparing global fits parameter estimations, *individual turnover* rates and the *delta ages* we grouped all samples together and modelled them collectively. Again, we found that the same scenario 5 best fitted the ^14^C results with a *renewing fraction* and *turnover rate* estimate of 0.64 and 3.3% respectively (Supplementary Fig. 5d, supplementary Table 4).

### Reduced neurogenesis throughout life is not a likely mechanism of addiction vulnerability

To investigate whether a reduced neurogenesis throughout life could be a common mechanism of addiction vulnerability in our two addiction groups, we compared the cell turnover between the healthy control group and the alcohol and cocaine groups. The same scenarios used for modelling non-neuronal samples were applied for the neuronal samples but also more complex models in order to better describe changes in adult neurogenesis over time and potential selective death of cells as observed in stereology data. In total, 11 different models were tested (Supplementary Table 2). However, we found that all global fits allowing a *renewing fraction* resulted in unrealistically high *turnover rates* (Supplementary Table 5). This is due to some samples before the rise of the atmospheric ^14^C curve can have two different solutions to their measured ^14^C values (Supplementary Fig. 7). Small differences in low *turnover rates* have larger effect on the modelled ^14^C value than changes in high *turnover rates*, which will penalize the small turnover solution when there is true variability. Thus, global model fitting was not optimal for this particular dataset and should be interpreted carefully but can be used for determining other parameters that do not have two solutions.

The best accepted model for all three groups was scenario 5, assuming a *renewing fraction* and a constant *turnover rate* (Supplementary Table 5, see Supplementary material). We thus modelled individual *turnover rates*, using the scenario 5 model and fixing the *renewing fraction* to the healthy control global fit estimation of 37% (Supplementary Fig. 8). The individual *turnover rates* did not correlate with age in any of the groups (Healthy control group: F[1, 22]= 25.00, R2= 0.05, p= 0.2822; Alcohol group: F[1, 19]= 13.39, R2= 0.13, p= 0.1093; Cocaine group: F[1, 15]= 0.98, R2=0.20, p=0.0686, linear regression, see Supplementary Fig. 9a), nor was there a significant difference between the healthy control group and the alcohol or cocaine groups (Fig. 3a, p> 0.05, Mann-Whitney U test). The average cell ages were modelled, using the individual *turnover rates* and the *delta age* was calculated for each sample. There was no significant correlation with neuronal *delta ages* and age (Healthy control group: F[1, 22]= 0.59, R2= 0.03, p= 0.4519; Alcohol group: F[1, 19]= 1.04, R2= 0.05, p= 0.3203; Cocaine group: F[1, 15]= 0.49, R2= 0.03, p= 0.4936, linear regression, see Supplementary Fig. 9b), nor was there significant difference when comparing the *delta ages* across the three groups (Fig. 3b, p> 0.05, Mann-Whitney U test).

**Figure 3:**
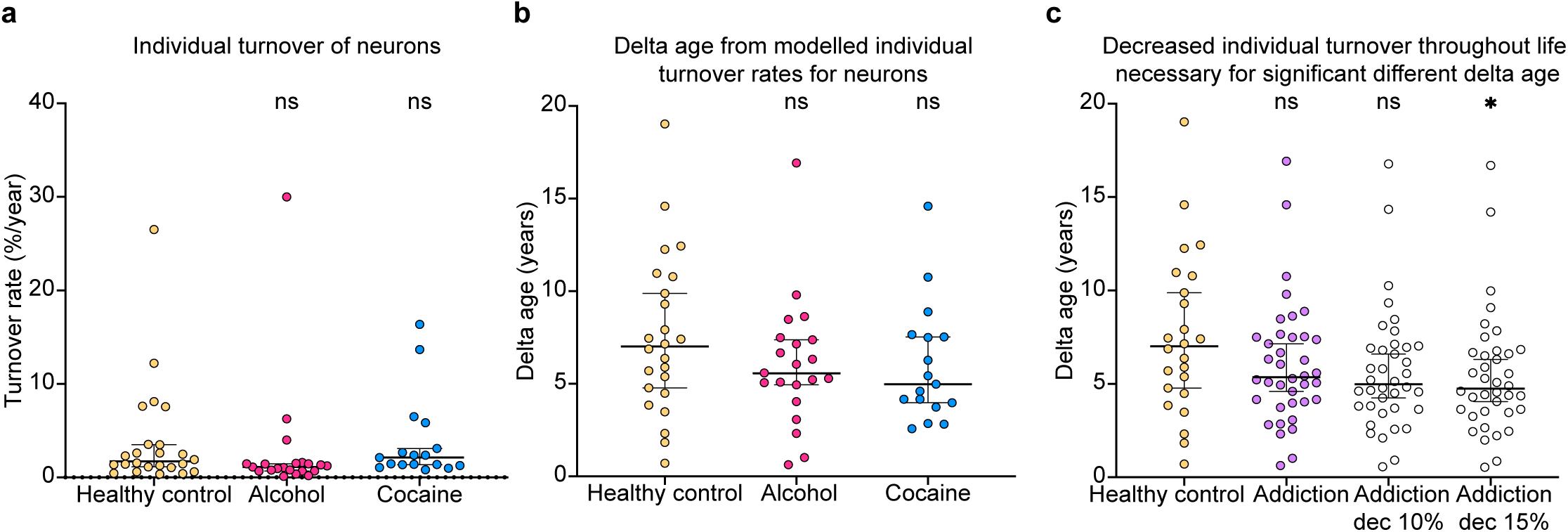
Individual modelling of neurons. **(a)** Comparison of annual turnover rates of neurons. There was no significant difference between healthy controls and alcohol or cocaine groups when comparing the annual turnover rates. **(b** and **c)** Comparison of neuronal delta age. **(b)**There was no significant difference when comparing healthy controls with the alcohol or cocaine group. **(c)** A reduction of 15% resulted in a significant difference between the abuse groups (cocaine and alcohol together) and healthy controls (p=0.0330). All tests were performed with Mann-Whitney U test, ns= non-significant (p> 0.05). Error bars show median with 95% confidence interval. Delta age= the difference in the person’s chronological age and the average cell age in years.

If there is a common mechanism of addiction between the alcohol and cocaine groups these groups could be studied together. Therefore we combined them in an “addiction group”. We could not detect any significant difference in neuronal *delta age* between the healthy control group and the addiction group (Fig. 3c, p= 0.1643, Mann-Whitney U test). To investigate how much reduction in turnover would be required for us to detect a significant difference, we modelled the average cell age after reducing the individual *turnover rates* with 10 and 15% and compared *delta ages*. We found that a significant difference could be detected in after a 15% reduction of the individual *turnover rates* throughout life in the addiction group, but not a reduction of 10% (Fig. 3c, 10% reduction: p= 0.0570; 15% reduction: p=0.0289, Mann-Whitney U test). The median individual *turnover rate* of the healthy controls is 1.7% and the median of the addiction group without any reduction is 1.4%. The median individual *turnover rate* after a 15% reduction within the addiction is 1%, thus up to 40% decrease in individual *turnover rates* over a lifetime cannot be excluded with our sample size.

### Significant adult neurogenesis persists after decades of alcohol abuse, but biologically relevant changes cannot be excluded

In rodent models, alcohol and cocaine affects the generation of new neurons^18^. Models with changes over a shorter period of time will be much more difficult to explore without a robust global model, which was not possible with this dataset. While we did not have data on disease duration of our cocaine group, we had information on years of abuse for 18 out of 23 individuals in the alcohol group. If there is a reduction of neurogenesis dependent on a continuous effect of reduced neurogenesis with alcohol use, one would expect a correlation between cell turnover and years of abuse. No significant correlation was found between years of abuse and neuronal *delta age* in the alcohol group (Fig. 4a, r= -0.3028, p= 0.2526, Spearman correlation). The years of abuse for most individuals was very long, representing a median of 30% of the individuals’ lifetime and median of 20 years of abuse. We modelled the *delta age* for the alcohol group if the *individual turnover* was reduced to 20 or 30% the last 20 years and found that we could detect a significant difference with a reduced neurogenesis of 30% for the alcohol group (Fig. 4b, 20% reduction: p= 0.0859; 30% reduction: p= 0.0365, Mann-Whitney U test).

**Figure 4:**
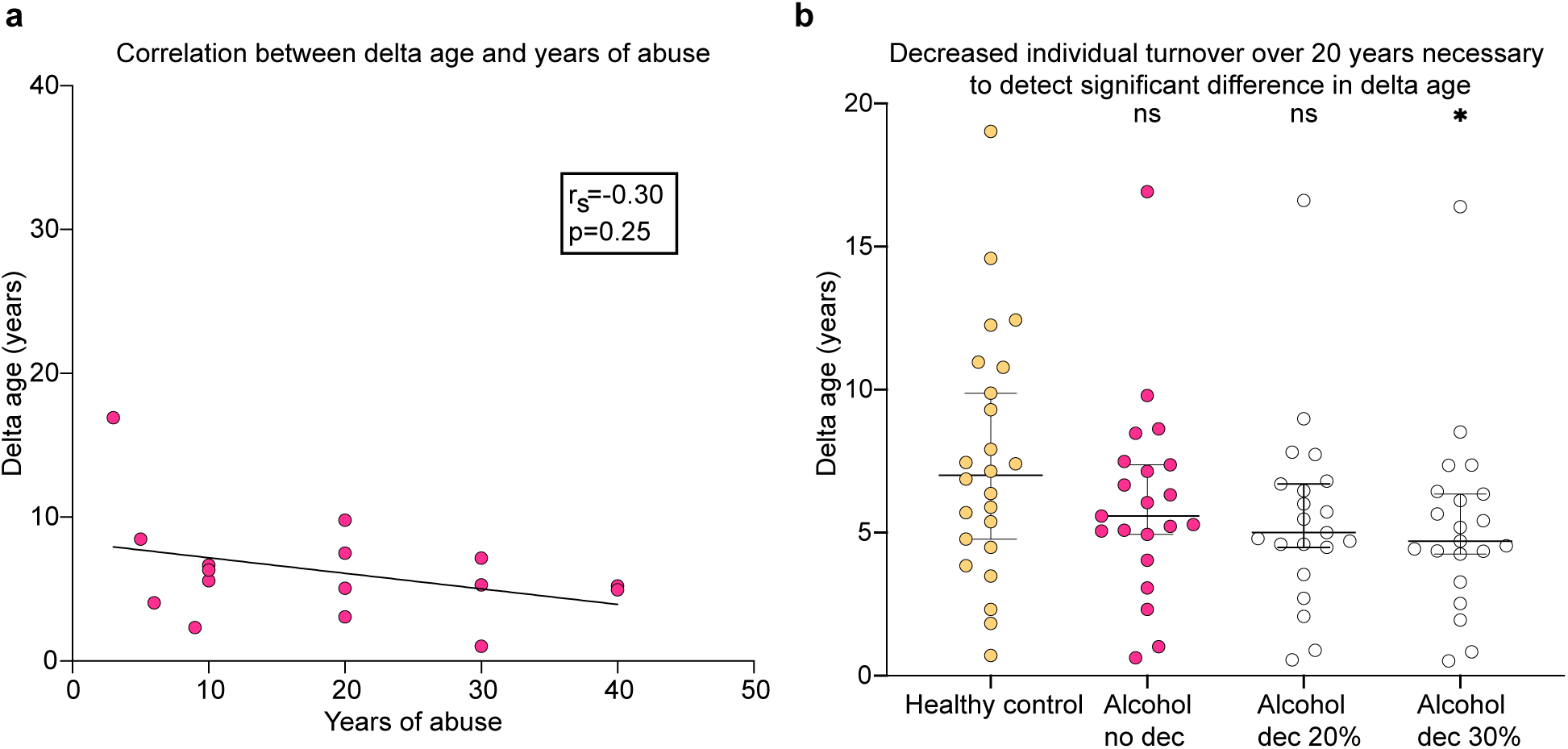
Investigation of reduced neuronal turnover with years of alcohol abuse. (**a**) Correlation between delta age of neurons and disease duration. There was no significant correlation with years of abuse for the alcohol group (r_s_=Spearman correlation coefficient, p values were considered non-significant if p> 0.05). **(b)** Comparison of neuronal delta age if reducing individual turnover rates 20 years before death. A significant difference (p= 0.0365) in delta age could be found after reducing individual turnover rates with 30% during 20 years before death. Tests performed with Mann-Whitney U test, ns= non-significant, (p > 0.05). Error bars show median with 95% confidence interval. Delta age= the difference in the person’s chronological age and the average cell age in years.

Theoretically, a reduction in adult neurogenesis could be masked by selective loss of old neurons and would lead to a less sensitive analysis. One stereology study have found that relatively more CA neurons than DG neurons die in younger individuals with alcohol abuse, but no difference in older subjects^29^. Other studies have failed to reproduce these results^43^. We modelled the *delta age* after a selective death of 25% of CA neurons (based on the lifetime loss in healthy controls) and tested what the minimum reduced individual *turnover rates* would have to be for us to detect a difference (Supplementary Fig. 10). To detect a significant difference in neuronal *delta age* a 40% (p= 0.0153, Mann-Whitney U test) reduction in individual *turnover rates* throughout life and 60% (p= 0271, Mann-Whitney U test) reduction after disease onset would be required.

## Discussion

Here we demonstrate that there is significant turnover of neuronal and non-neuronal cells in the hippocampus of individuals with long-term alcohol or cocaine use comparable to healthy control. The substantial variability in cell turnover within all three group limits the possibility to detect differences with our sample size. This variability could partly be explained by variability in adult neurogenesis. In rodents, genetics, physical activity and living in an enriched environment have substantial influence on the extent of adult neurogenesis^44–46^ and these factors may vary greatly within our healthy control group. It is likely that the variability of average age observed also largely influenced by differences in CA to DG proportions as seen in stereology data^38–42^.

Assuming that a substantial reduction of more than 50% in hippocampal neuronal *turnover rate* is required to impact on cognitive disturbances^18^, our data argues against the theory of reduced neurogenesis as a mechanism of addiction vulnerability. However, it is important to note that we would not be able to exclude up to a 40% reduction of individual *turnover rates* from birth. While detecting alterations in neurogenesis over shorter periods of time is not possible with the carbon dating strategy, we can exclude scenarios of a complete shut off neurogenesis after severe alcohol abuse. Even individuals with over 20 years severe alcohol use, representing more than a third of their lives, demonstrate significant neurogenesis after onset of abuse.

Studies of adult neurogenesis after alcohol exposure have reported various results, however many demonstrate a reduction of around 40-50%^18^, which would be just at the detection limit in this study. In addition, abstinence following alcohol dependence increases neurogenesis in rodent models and many individuals within this study may have been periodic drinkers, and many had records of hospitalization for abstinence^23^. Thus, we cannot exclude smaller differences after alcohol or cocaine exposure.

Potential masking of reduced neurogenesis by selective loss of old neurons might further reduce the sensitivity of detecting reduced neurogenesis using carbon dating. Although most likely not a realistic scenario, it is important to note that we can exclude scenarios of shut off adult neurogenesis even at a selective cell loss of 25% of prenatal neurons. In conclusion, we cannot exclude dynamic changes after use of alcohol or cocaine, however our data suggests that the overall turnover remain similar in healthy controls and individuals with alcohol or cocaine abuse, suggesting that reduced adult neurogenesis is not an underlying mechanism of addiction vulnerability.

---

**Supplementary Figure 1:**
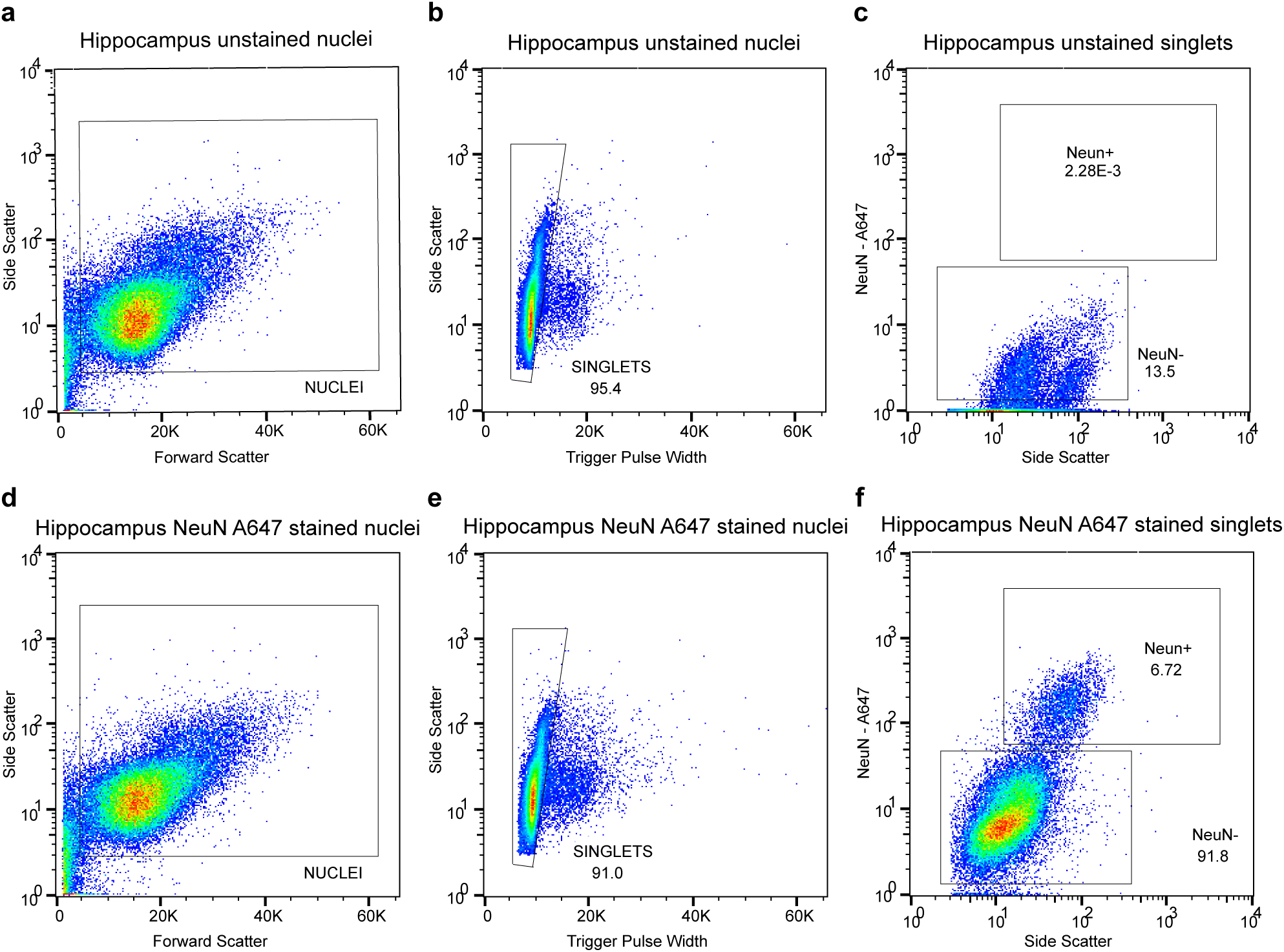
Gating strategy for sorting of neuronal and non-neuronal population. Panels show gating for **(a-c)** unstained nuclei and **(d-f)** stained nuclei using NeuN antibody conjugated to Alexa 647 (A647) fluorophore. **(a** and **d)** Whole nuclei are selected then (**b** and **e**) doublet nuclei are filtered out and only singlets are selected and finally **(c** and **f)** neurons are selected through fluorescent activity of the NeuN antibody.

**Supplementary Figure 2:**
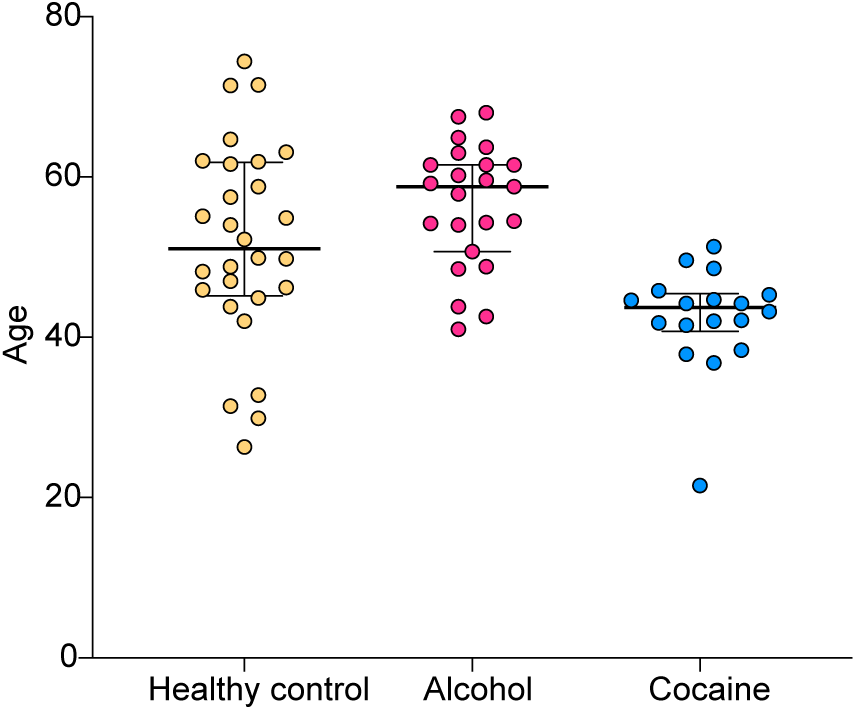
Ages of healthy control, alcohol and cocaine group. Error bars shows median with interquartile range.

**Supplementary Figure 3:**
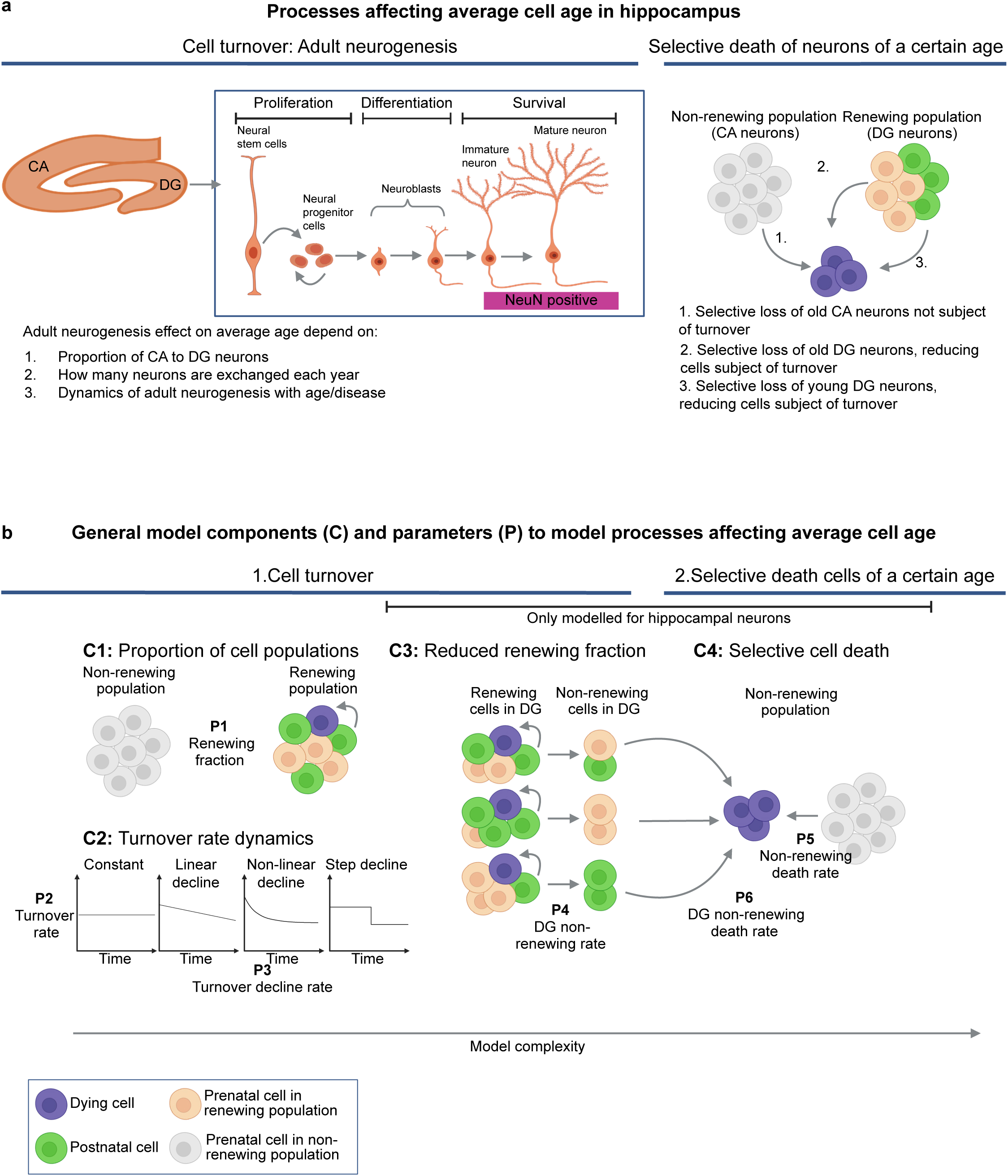
Schematic figures of the main concepts that affect average cell age that can be modelled: cell turnover and selective cell loss. **(a)** Schematic figure of processes within the hippocampus that affect neuronal average age. We collect nuclei from neurons positive for the neuronal marker NeuN, which is expressed in cornu ammonis (CA) neurons and more mature neurons in the dentate gyrus (DG). The effect of cell turnover (adult neurogenesis) on neuronal average cell age is dependent on 1. The proportion of CA to DG neurons 2. How many neurons are exchanged each year. 3. The dynamics of adult neurogenesis with age or disease. **(b)** Schematic figure of the general model components (C) and parameters (P) that make up the scenarios of processes affecting average cell age. Up to 4 components (C1 – C4) and 6 parameters (P1-P6) are used to model processes that may affect the average age of a population. C1 describes the non-renewing and renewing cell populations with the parameter (P1) *renewing fraction,* which is the proportion of the renewing population. C2 describes the turnover rate dynamics with age and entails two parameters: (P2) Turnover rate, which is the percent of cells within the *renewing fraction* that are exchanged each year, and (P3) turnover decline rate which is a parameter that describes a decrease of turnover with a specific function. C3 describes a more complex scenario, only for neurons where the *renewing fraction* (the DG) can decline with age as cells go from a renewing state to a non-renewing state by parameter (P4) DG non-renewing rate. C4 describes selective cell loss of cells within the CA with the parameter (P5) non-renewing death rate or within the DG with the parameter (P6) DG non-renewing death rate. In total 8 scenarios, used for modelling of both non-neuronal and neuronal populations scenarios were composed of components C1 and C2 with increasing complexity, defined as number of parameters. For the neuronal population we developed 3 highly complex scenarios composing of all four components and parameters (see Supplementary Table 2). Created with Biorender.com.

**Supplementary Figure 4:**
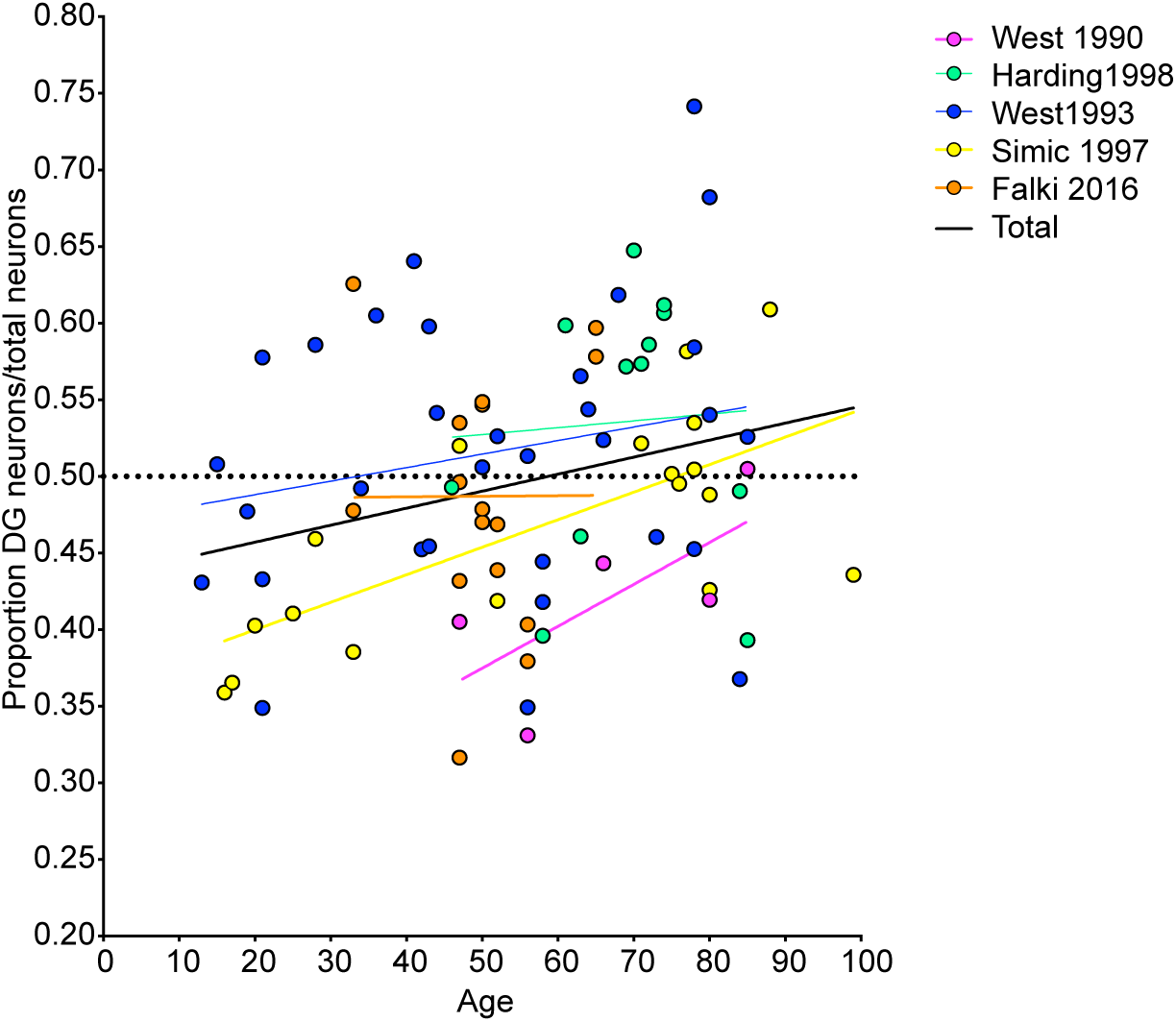
Proportion DG neurons of total neurons from Dentate gyrus (DG) and cornu ammonis (CA) regions from 5 stereology studies^38–42^.

**Supplementary Figure 5:**
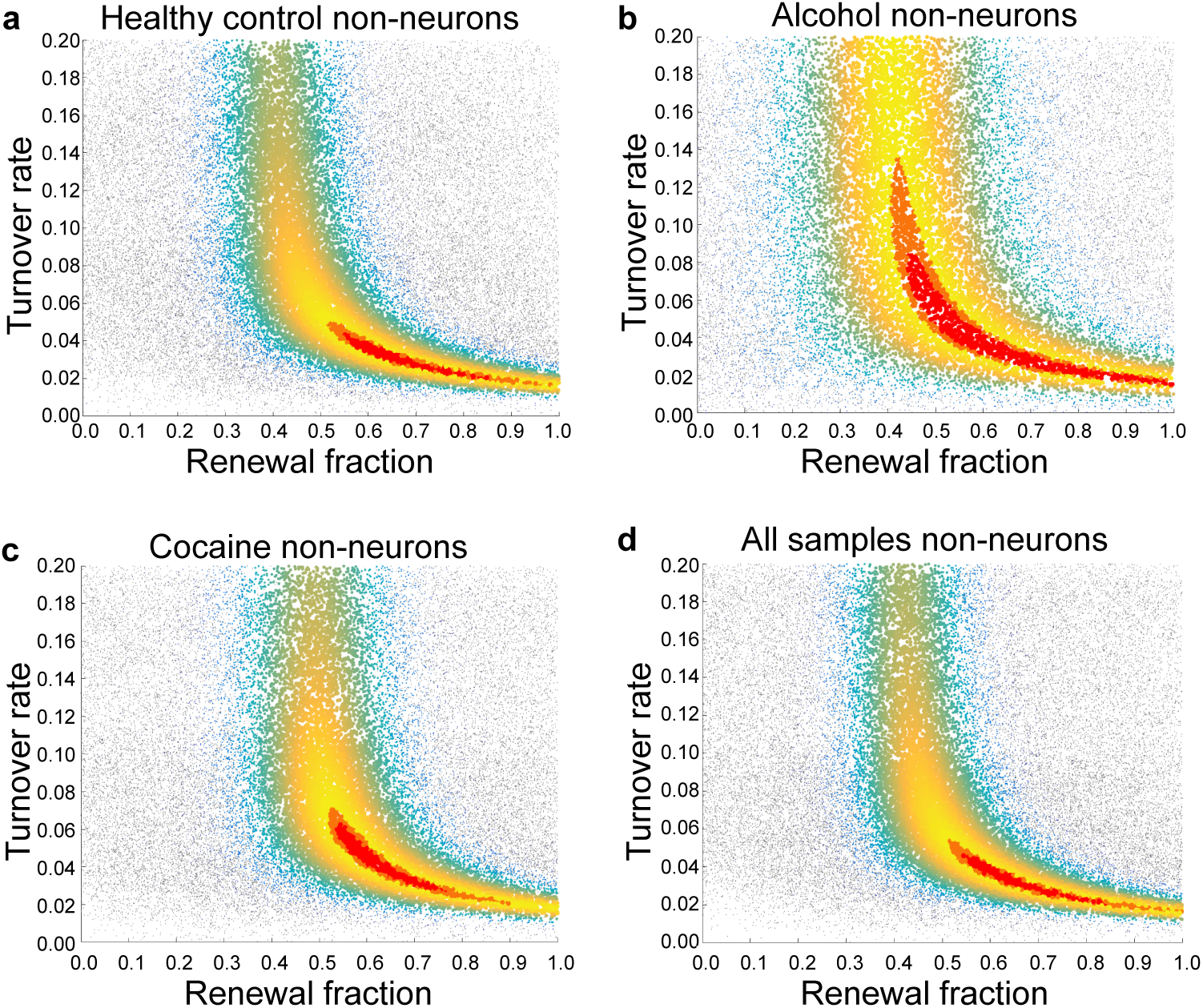
Markov Chain Monte Carlo simulations of scenario 5 for non-neuron samples in the healthy control, alcohol and cocaine groups. Scenario 5 estimates two parameters: 1. Renewal fraction (x axis) and turnover rate (y axis). The red area is the parameter estimates with highest likelihood. Models were tested on **(a)** healthy control group, **(b)** alcohol group and **(c)** cocaine group respectively. **(d)** Shows simulations of all samples together.

**Supplementary Figure 6:**
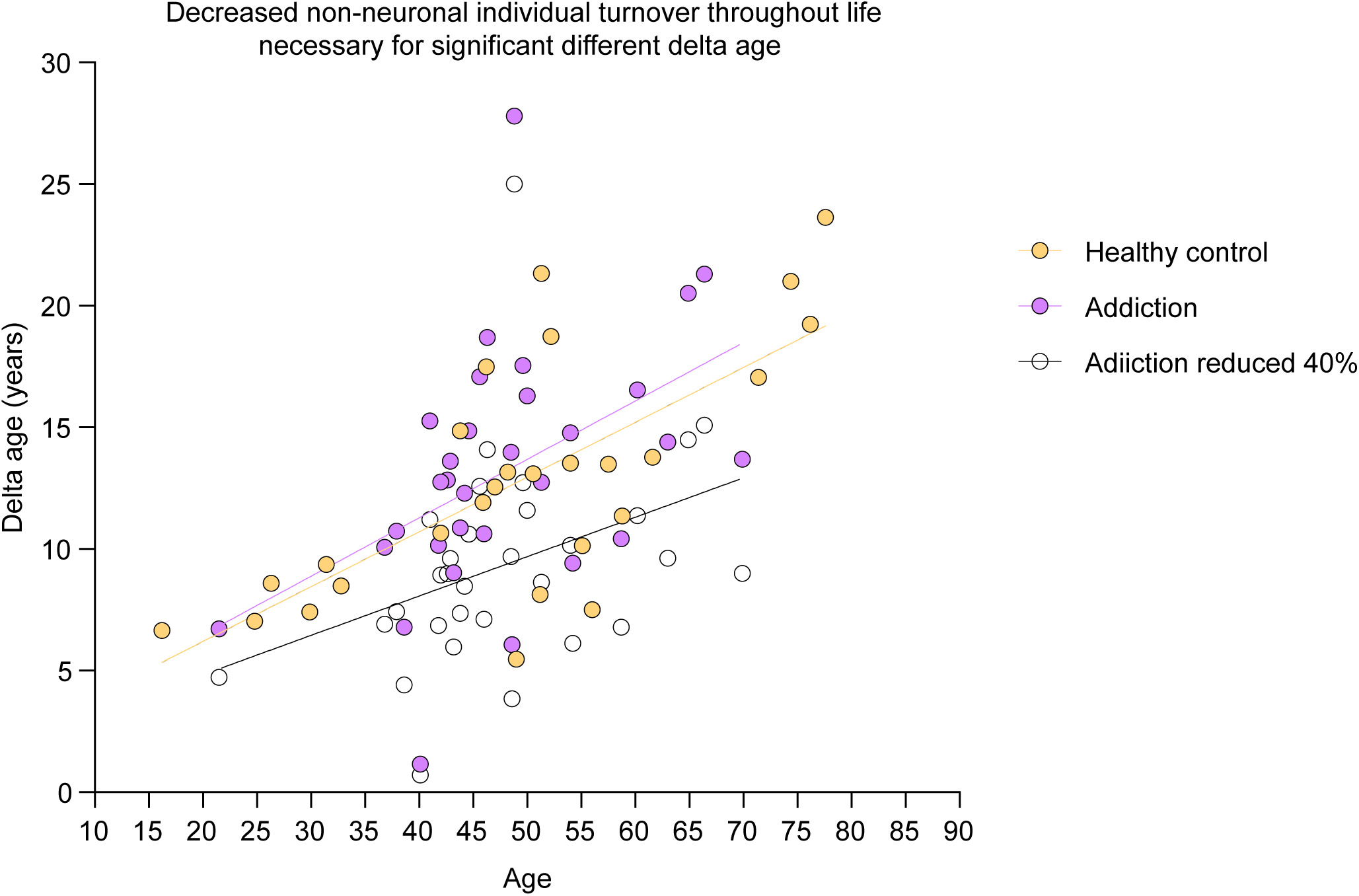
Required decrease of individual turnover rates of non-neurons for significant different delta age. There was no significant difference between healthy controls and the addiction group delta ages (alcohol and cocaine together) (F[2, 56]= 305.4, R2= 0.92, p < 0.0001. Age was significant: F[1, 56]= 280.8, p< 0.0001. Group effect was not significant: F[1, 56]= 0.6353, p= 0.4288, multiple linear regression). A 40% decrease in non-neuronal individual turnover rates was required for a significant difference in delta age (F[2, 56]= 256.1, R2= 0.90, p < 0.0001. Age was significant: F[1, 56]= 322.4, p< 0.0001. Group effect: F[1, 56]= 9.817, p= 0.0028, multiple linear regression).

**Supplementary Figure 7:**
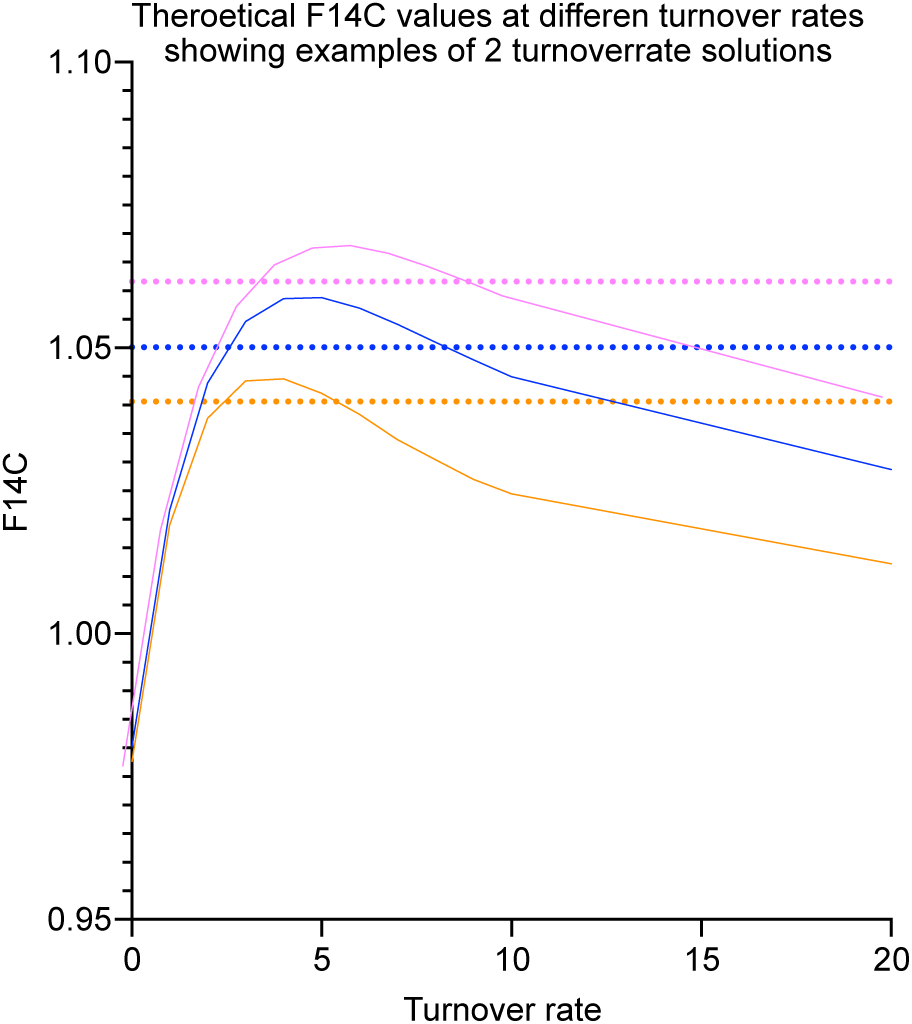
Theoretical examples of two turnover solutions. Two different turnover rates in some samples can result in the same ^14^C concentration. Y-axis represents the modelled F14C value and X axis represents different turnover rates modelled. The solid lines (purple, blue and orange) represent different individuals and the dashed lines represent their measured ^14^C values (color coded the same as the individual). The measured value intersects 2 different turnover rates, meaning that there are two solutions to the measured ^14^C value.

**Supplementary Figure 8:**
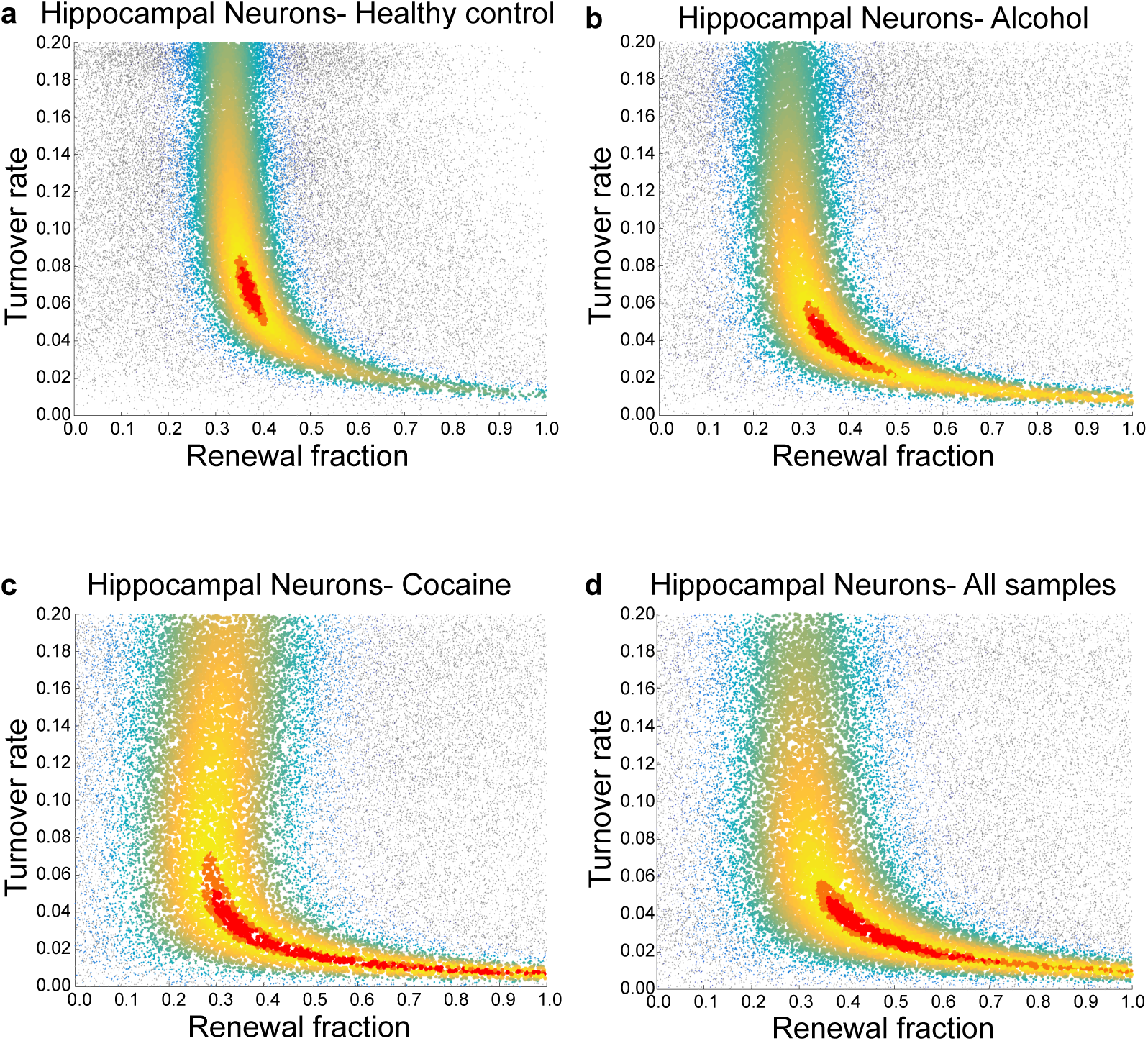
Markov Chain Monte Carlo simulations of scenario 5 for neuron samples. Scenario 5 estimates two parameters: 1. Renewal fraction (x axis) and turnover rate (y axis). The red area is the parameter estimates with highest likelihood. Models were tested on **(a)** healthy control group, **(b)** alcohol group and **(c)** cocaine group respectively. **(d)** Shows simulations of all samples together.

**Supplementary Figure 9:**
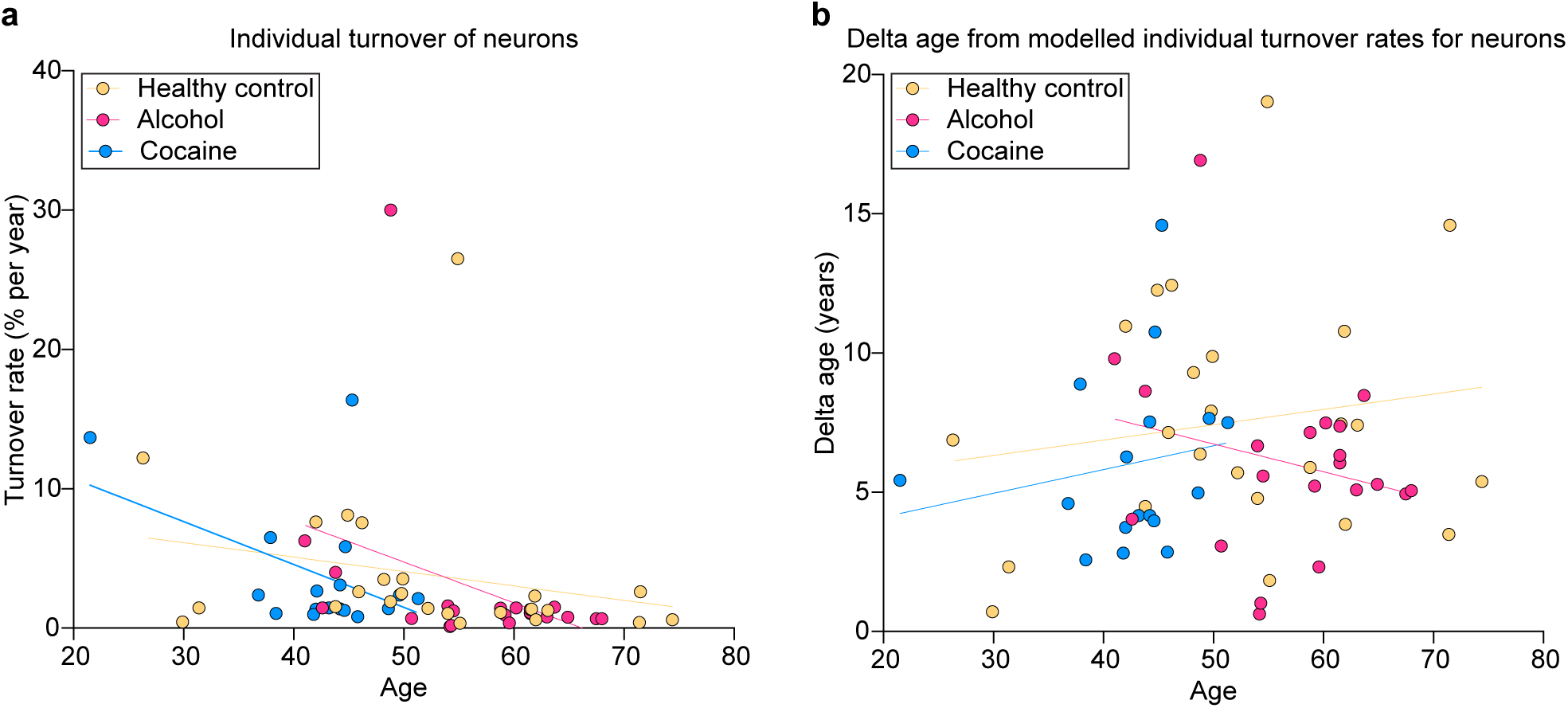
Correlation between neuronal turnover and age. **(a)** Correlation between annual turnover rate of neurons and age. The individual turnover rates did not correlate with age in any of the groups (Healthy control group: F[1, 22]= 25.00, R2= 0.05, p= 0.2822; Alcohol group: F[1, 19]= 13.39, R2= 0.13, p= 0.1093; Cocaine group: F[1, 15]= 0.98, R2= 0.20, p= 0.0686, linear regression. **(b)** Correlation between delta age for neurons and age. There was no significant correlation with delta age (Healthy control group: F[1, 22]= 0.59, R2= 0.03, p= 0.4519; Alcohol group: F[1, 19]= 1.04, R2= 0.05, p= 0.3203; Cocaine group: F[1, 15]= 0.49, R2= 0.03, p= 0.4936, linear regression). Delta age= the difference in the person’s chronological age and the average cell age in years.

**Supplementary Figure 10:**
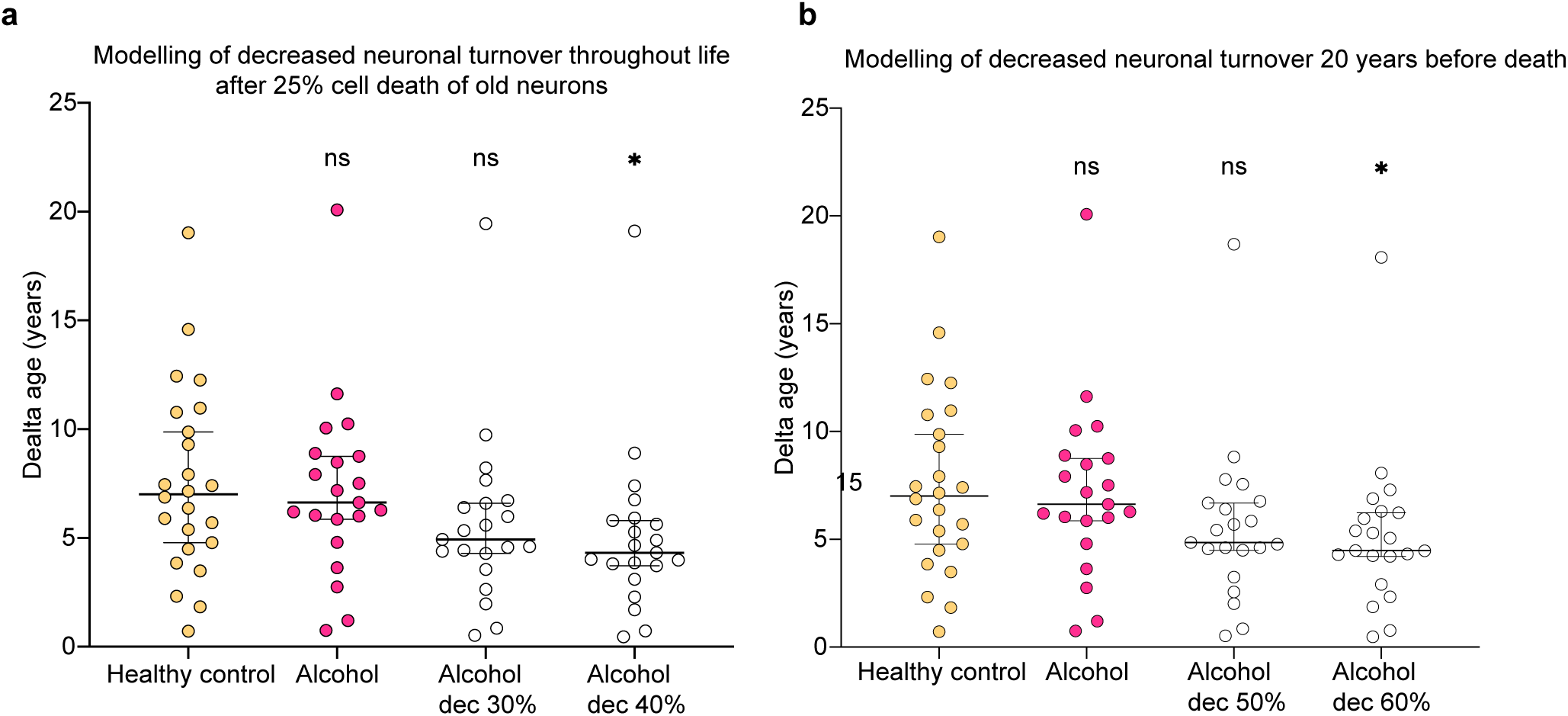
Individual turnover reduction necessary to detect a significant difference in person age-cell age difference if selective cell loss of old neurons. Comparison of delta age if reducing individual turnover of neurons **(a)** during the whole life or **(b)** last 20 years of life. A significant difference could be found after reducing individual turnover of neurons with 40% (P=0.0153) during the whole life and at 70% (p= 0271) of the last 20 years. All tests were performed with Mann-Whitney U test, ns= non-significant (p> 0.05). Error bars show median with 95% confidence interval. Delta age= the difference in the person’s chronological age and the average cell age in years.

**Supplementary Table 1.**
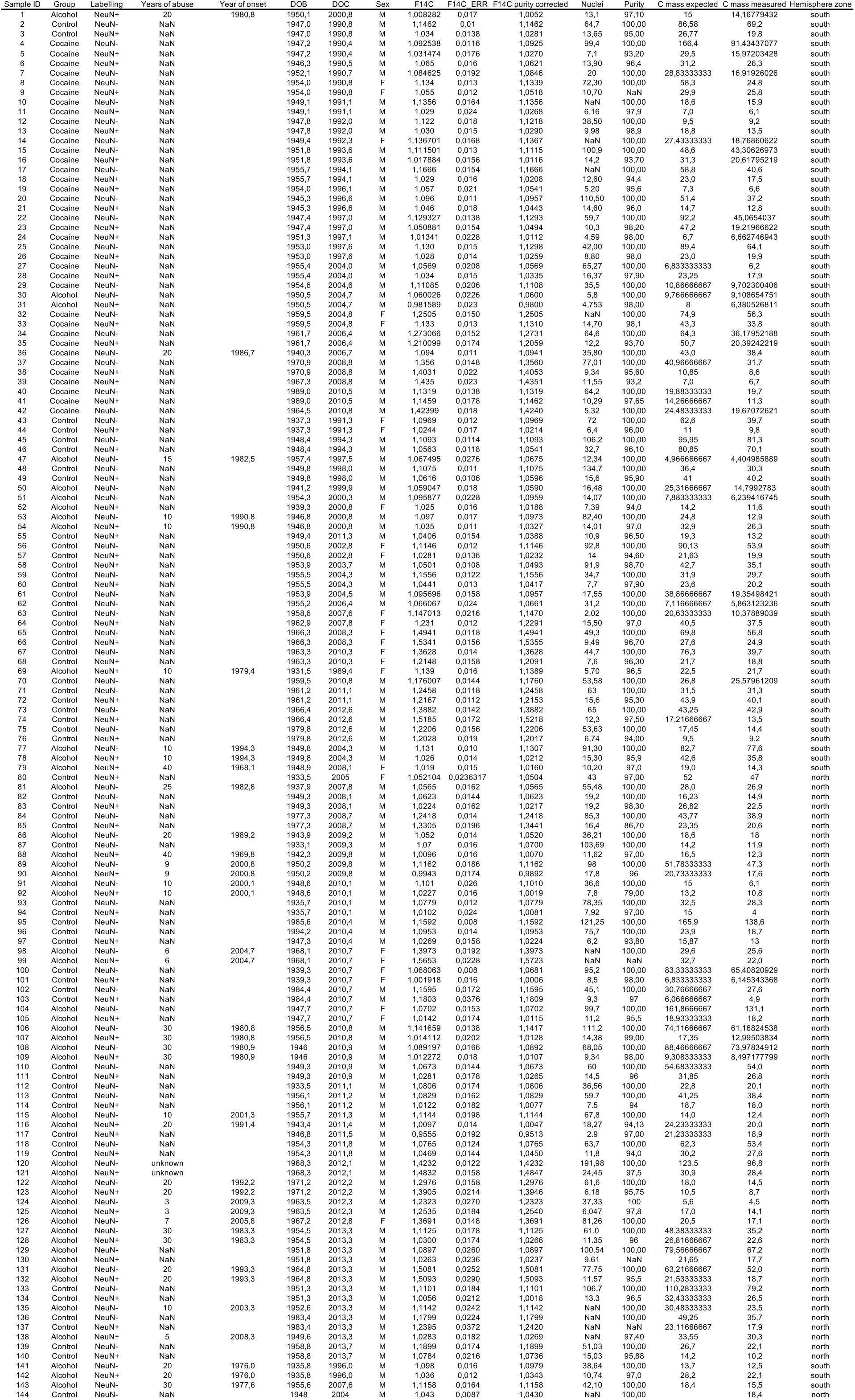

**Supplementary Table 2.**
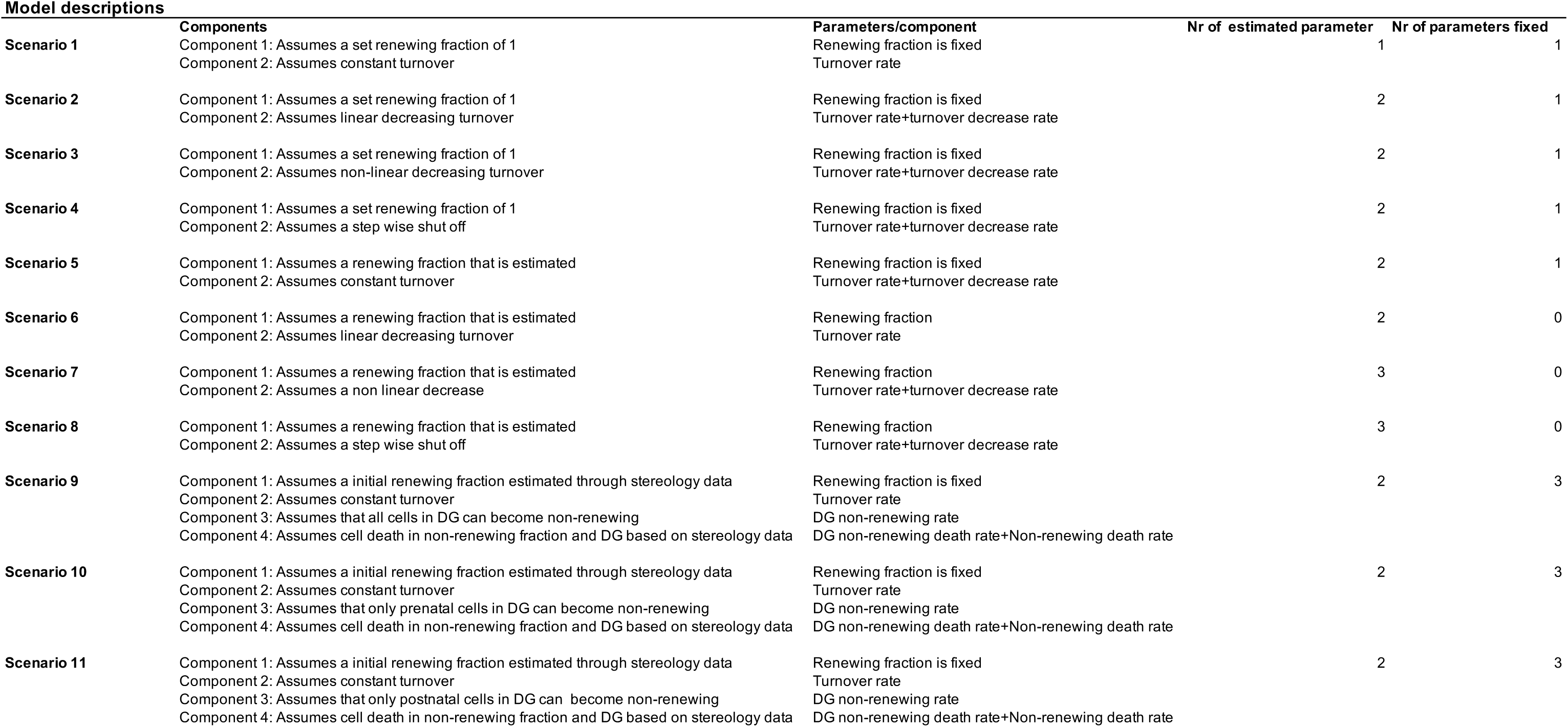

**Supplementary Table 3.**

| Weights non-neuronal samples |  |  |  |  |  | Weights Neuronal samples |  |  |  |  |  |
| --- | --- | --- | --- | --- | --- | --- | --- | --- | --- | --- | --- |
| Sample ID | Lag 0 | Lag 0.5 | Lag 1 | Lag 1.5 | Lag 2 | Sample ID | Lag 0 | Lag 0.5 | Lag 1 | Lag 1.5 | Lag 2 |
| 2 | 0,4335 | 0,4470 | 0,4587 | 0,4674 | 0,4748 | 1 | 0,138388 | 0,143881 | 0,147651 | 0,148889 | 0,149634 |
| 4 | 0,3820 | 0,3926 | 0,4020 | 0,4090 | 0,4159 | 3 | 0,246596 | 0,256927 | 0,26495 | 0,270189 | 0,27509 |
| 7 | 0,1697 | 0,1754 | 0,1804 | 0,1838 | 0,1870 | 5 | 0,19779 | 0,205456 | 0,211379 | 0,215235 | 0,219326 |
| 8 | 0,3391 | 0,3505 | 0,3599 | 0,3680 | 0,3752 | 6 | 0,282185 | 0,293121 | 0,301702 | 0,30711 | 0,312332 |
| 10 | 0,1974 | 0,2021 | 0,2065 | 0,2108 | 0,2140 | 9 | 0,385328 | 0,40233 | 0,415223 | 0,424994 | 0,433993 |
| 12 | 0,1121 | 0,1141 | 0,1169 | 0,1201 | 0,1229 | 11 | 0,094131 | 0,097292 | 0,099925 | 0,102057 | 0,103826 |
| 14 | 0,1851 | 0,1872 | 0,1898 | 0,1939 | 0,1982 | 13 | 0,209855 | 0,2158 | 0,221986 | 0,22837 | 0,234201 |
| 15 | 0,2999 | 0,3075 | 0,3134 | 0,3190 | 0,3244 | 16 | 0,260276 | 0,269513 | 0,27567 | 0,280934 | 0,286291 |
| 17 | 0,1940 | 0,2199 | 0,2457 | 0,2549 | 0,2632 | 18 | 0,14485 | 0,166024 | 0,186749 | 0,194029 | 0,200782 |
| 20 | 0,3103 | 0,3173 | 0,3247 | 0,3305 | 0,3344 | 19 | 0,086272 | 0,089563 | 0,092012 | 0,093726 | 0,095675 |
| 22 | 0,2471 | 0,2517 | 0,2580 | 0,2642 | 0,2684 | 21 | 0,1517 | 0,156568 | 0,160906 | 0,163871 | 0,16591 |
| 25 | 0,2269 | 0,2315 | 0,2366 | 0,2427 | 0,2485 | 23 | 0,171889 | 0,176652 | 0,181874 | 0,186408 | 0,189593 |
| 27 | 0,0473 | 0,0549 | 0,0628 | 0,0652 | 0,0675 | 24 | 0,077982 | 0,079929 | 0,081788 | 0,084163 | 0,086129 |
| 29 | 0,0578 | 0,0604 | 0,0629 | 0,0643 | 0,0660 | 26 | 0,186177 | 0,191711 | 0,196638 | 0,201891 | 0,207289 |
| 30 | 0,0565 | 0,0583 | 0,0601 | 0,0605 | 0,0613 | 28 | 0,100265 | 0,117343 | 0,135019 | 0,140377 | 0,145371 |
| 32 | 0,2721 | 0,1998 | 0,1224 | 0,0545 | 0,0185 | 31 | 0,056727 | 0,059043 | 0,061046 | 0,061437 | 0,018647 |
| 34 | 0,2940 | 0,2783 | 0,2605 | 0,2617 | 0,2636 | 33 | 0,313246 | 0,230042 | 0,14234 | 0,064 | 0,02191 |
| 36 | 0,2060 | 0,2054 | 0,2050 | 0,2109 | 0,2171 | 35 | 0,254191 | 0,238168 | 0,221546 | 0,222522 | 0,224267 |
| 37 | 0,5704 | 0,5787 | 0,5880 | 0,5952 | 0,6015 | 38 | 0,191618 | 0,193468 | 0,195968 | 0,198111 | 0,199724 |
| 40 | 0,1770 | 0,1775 | 0,1817 | 0,1857 | 0,1879 | 39 | 0,20885 | 0,212685 | 0,215673 | 0,220504 | 0,226592 |
| 42 | 0,4545 | 0,4570 | 0,4243 | 0,3586 | 0,2683 | 41 | 0,136516 | 0,136575 | 0,139571 | 0,142561 | 0,144099 |
| 43 | 0,3434 | 0,3526 | 0,3625 | 0,3706 | 0,3775 | 44 | 0,126657 | 0,131312 | 0,135742 | 0,13883 | 0,142234 |
| 45 | 0,3312 | 0,3381 | 0,3460 | 0,3533 | 0,3582 | 46 | 0,33277 | 0,343009 | 0,352513 | 0,360187 | 0,36581 |
| 47 | 0,0099 | 0,0142 | 0,0185 | 0,0230 | 0,0275 | 49 | 0,327958 | 0,335333 | 0,341045 | 0,344561 | 0,3505 |
| 48 | 0,3060 | 0,3100 | 0,3139 | 0,3170 | 0,3221 | 52 | 0,138528 | 0,143696 | 0,147241 | 0,149505 | 0,151264 |
| 50 | 0,1302 | 0,1323 | 0,1337 | 0,1340 | 0,1340 | 54 | 0,276867 | 0,287908 | 0,29592 | 0,300208 | 0,303755 |
| 51 | 0,0658 | 0,0679 | 0,0695 | 0,0704 | 0,0712 | 55 | 0,096016 | 0,096865 | 0,097672 | 0,100937 | 0,103119 |
| 53 | 0,1361 | 0,1404 | 0,1437 | 0,1458 | 0,1474 | 57 | 0,155759 | 0,163022 | 0,168699 | 0,170874 | 0,173618 |
| 56 | 0,2296 | 0,2383 | 0,2458 | 0,2484 | 0,2526 | 58 | 0,24291 | 0,25066 | 0,259451 | 0,268892 | 0,276559 |
| 59 | 0,1545 | 0,1825 | 0,2102 | 0,2185 | 0,2270 | 60 | 0,147502 | 0,175953 | 0,203804 | 0,211964 | 0,220532 |
| 61 | 0,1186 | 0,1212 | 0,1237 | 0,1280 | 0,1324 | 64 | 0,689036 | 0,571344 | 0,429663 | 0,378933 | 0,339811 |
| 62 | 0,0379 | 0,0432 | 0,0490 | 0,0509 | 0,0526 | 66 | 0,642827 | 0,654217 | 0,669133 | 0,688419 | 0,689432 |
| 63 | 0,0876 | 0,0514 | 0,0143 | 0,0044 | 0,0234 | 68 | 0,459361 | 0,387511 | 0,295822 | 0,246216 | 0,205316 |
| 65 | 0,8478 | 0,8668 | 0,8892 | 0,9158 | 0,9203 | 69 | 0,292032 | 0,300938 | 0,308389 | 0,314818 | 0,318864 |
| 67 | 0,6900 | 0,5865 | 0,4497 | 0,3747 | 0,3145 | 72 | 0,393571 | 0,393255 | 0,391926 | 0,366905 | 0,344357 |
| 70 | 0,3234 | 0,2571 | 0,1749 | 0,1048 | 0,0329 | 74 | 0,443313 | 0,449556 | 0,45986 | 0,473576 | 0,475158 |
| 71 | 0,3755 | 0,3777 | 0,3774 | 0,3552 | 0,3350 | 76 | 0,148842 | 0,153642 | 0,158801 | 0,161161 | 0,162662 |
| 73 | 0,7131 | 0,7260 | 0,7449 | 0,7677 | 0,7729 | 78 | 0,195743 | 0,200049 | 0,203262 | 0,206734 | 0,210967 |
| 75 | 0,2731 | 0,2829 | 0,2930 | 0,2977 | 0,3009 | 79 | 0,115453 | 0,118672 | 0,121152 | 0,123585 | 0,125015 |
| 77 | 0,2640 | 0,2679 | 0,2712 | 0,2758 | 0,2813 | 80 | 0,097601 | 0,100683 | 0,103565 | 0,106695 | 0,109154 |
| 81 | 0,1257 | 0,1305 | 0,1339 | 0,1331 | 0,1339 | 83 | 0,137071 | 0,139968 | 0,143681 | 0,145133 | 0,14349 |
| 82 | 0,1139 | 0,1154 | 0,1181 | 0,1193 | 0,1180 | 85 | 0,320743 | 0,324262 | 0,328051 | 0,332579 | 0,339143 |
| 84 | 0,4511 | 0,4579 | 0,4644 | 0,4712 | 0,4814 | 88 | 0,091929 | 0,094299 | 0,096084 | 0,099042 | 0,101516 |
| 86 | 0,1097 | 0,1103 | 0,1127 | 0,1146 | 0,1181 | 90 | 0,089874 | 0,091247 | 0,092577 | 0,094052 | 0,09534 |
| 87 | 0,0873 | 0,0883 | 0,0913 | 0,0935 | 0,0971 | 92 | 0,093305 | 0,095003 | 0,096624 | 0,098665 | 0,101748 |
| 89 | 0,1104 | 0,1115 | 0,1128 | 0,1146 | 0,1162 | 94 | 0,019097 | 0,019556 | 0,020116 | 0,020419 | 0,021071 |
| 91 | 0,0378 | 0,0382 | 0,0388 | 0,0396 | 0,0408 | 97 | 0,091804 | 0,095105 | 0,097191 | 0,0992 | 0,102293 |
| 93 | 0,1509 | 0,1536 | 0,1575 | 0,1601 | 0,1650 | 99 | 0,411074 | 0,420834 | 0,430623 | 0,441772 | 0,452414 |
| 95 | 0,4974 | 0,5031 | 0,5100 | 0,5245 | 0,5371 | 101 | 0,056867 | 0,057797 | 0,059282 | 0,060609 | 0,061369 |
| 96 | 0,1143 | 0,1157 | 0,1216 | 0,1243 | 0,1229 | 103 | 0,027904 | 0,028572 | 0,02925 | 0,029953 | 0,030617 |
| 98 | 0,4874 | 0,5012 | 0,5144 | 0,5284 | 0,5424 | 105 | 0,082923 | 0,084588 | 0,086963 | 0,089225 | 0,090895 |
| 100 | 0,2250 | 0,2272 | 0,2323 | 0,2375 | 0,2405 | 107 | 0,019595 | 0,039415 | 0,059809 | 0,067869 | 0,076008 |
| 102 | 0,2453 | 0,2519 | 0,2584 | 0,2648 | 0,2710 | 109 | 0,053604 | 0,05473 | 0,056295 | 0,057233 | 0,05834 |
| 104 | 0,1243 | 0,1260 | 0,1292 | 0,1324 | 0,1350 | 111 | 0,111545 | 0,112307 | 0,113636 | 0,114531 | 0,11592 |
| 106 | 0,0381 | 0,0761 | 0,1147 | 0,1299 | 0,1453 | 114 | 0,037745 | 0,054037 | 0,070649 | 0,077835 | 0,084614 |
| 108 | 0,1150 | 0,1166 | 0,1196 | 0,1216 | 0,1239 | 116 | 0,102556 | 0,103254 | 0,102886 | 0,10553 | 0,107074 |
| 110 | 0,1363 | 0,1361 | 0,1374 | 0,1386 | 0,1404 | 117 | 0,073259 | 0,075617 | 0,076972 | 0,078556 | 0,079682 |
| 112 | 0,1007 | 0,1016 | 0,1029 | 0,1048 | 0,1074 | 119 | 0,127812 | 0,133613 | 0,13819 | 0,140904 | 0,145115 |
| 113 | 0,0563 | 0,0799 | 0,1039 | 0,1144 | 0,1242 | 121 | 0,58923 | 0,601949 | 0,61586 | 0,633019 | 0,649146 |
| 115 | 0,0488 | 0,0588 | 0,0690 | 0,0733 | 0,0770 | 123 | 0,201792 | 0,203151 | 0,205227 | 0,207062 | 0,208773 |
| 118 | 0,1470 | 0,1526 | 0,1573 | 0,1604 | 0,1652 | 125 | 0,427832 | 0,361978 | 0,273042 | 0,229546 | 0,177567 |
| 120 | 0,7617 | 0,7813 | 0,8018 | 0,8251 | 0,8484 | 128 | 0,084869 | 0,094964 | 0,105471 | 0,111188 | 0,114086 |
| 122 | 0,4098 | 0,4144 | 0,4200 | 0,4244 | 0,4290 | 130 | 0,050586 | 0,054844 | 0,05939 | 0,062038 | 0,063293 |
| 124 | 0,0967 | 0,0825 | 0,0626 | 0,0527 | 0,0409 | 132 | 0,290769 | 0,283022 | 0,275675 | 0,246161 | 0,208932 |
| 126 | 0,5041 | 0,5140 | 0,5253 | 0,5379 | 0,5513 | 134 | 0,075109 | 0,081413 | 0,088169 | 0,091906 | 0,093335 |
| 127 | 0,0824 | 0,0916 | 0,1014 | 0,1068 | 0,1096 | 137 | 0,097171 | 0,097234 | 0,097784 | 0,099787 | 0,102353 |
| 129 | 0,0608 | 0,0655 | 0,0706 | 0,0738 | 0,0752 | 138 | 0,087604 | 0,093352 | 0,09947 | 0,10334 | 0,104259 |
| 131 | 0,4427 | 0,4332 | 0,4238 | 0,3804 | 0,3242 | 140 | 0,125613 | 0,09241 | 0,055378 | 0,026721 | 0,002214 |
| 133 | 0,0859 | 0,0925 | 0,0998 | 0,1040 | 0,1054 | 142 | 0,286848 | 0,296721 | 0,303004 | 0,306766 | 0,311661 |
| 135 | 0,0651 | 0,0702 | 0,0758 | 0,0793 | 0,0807 |  |  |  |  |  |  |
| 136 | 0,2163 | 0,2170 | 0,2186 | 0,2232 | 0,2293 |  |  |  |  |  |  |
| 139 | 0,2125 | 0,1562 | 0,0928 | 0,0445 | 0,0037 |  |  |  |  |  |  |
| 141 | 0,1629 | 0,1670 | 0,1699 | 0,1719 | 0,1744 |  |  |  |  |  |  |
| 143 | 0,0646 | 0,0827 | 0,0990 | 0,1033 | 0,1079 |  |  |  |  |  |  |
| 144 | 0,2177 | 0,2236 | 0,2285 | 0,2361 | 0,2442 |  |  |  |  |  |  |

**Supplementary Table 4.**
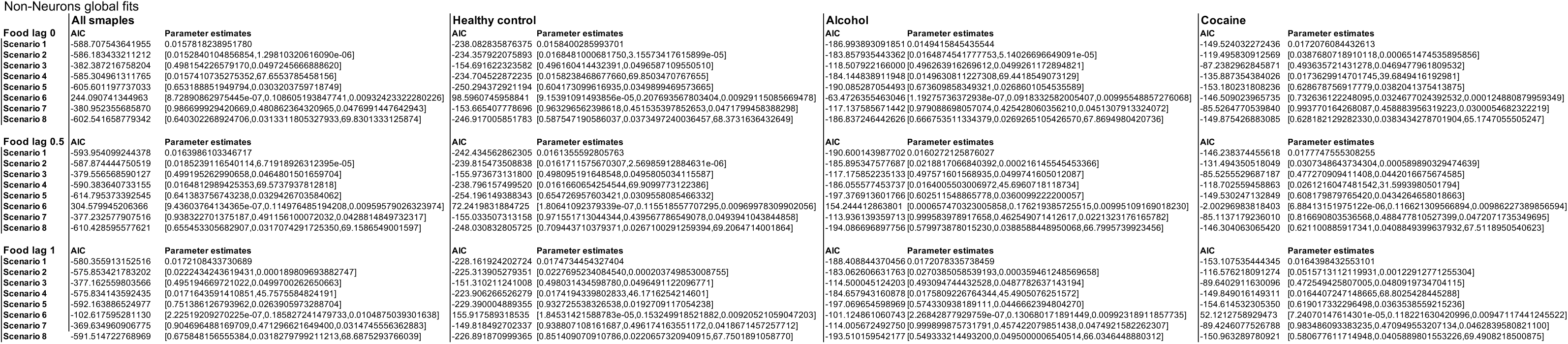

**Supplementary Table 5.**
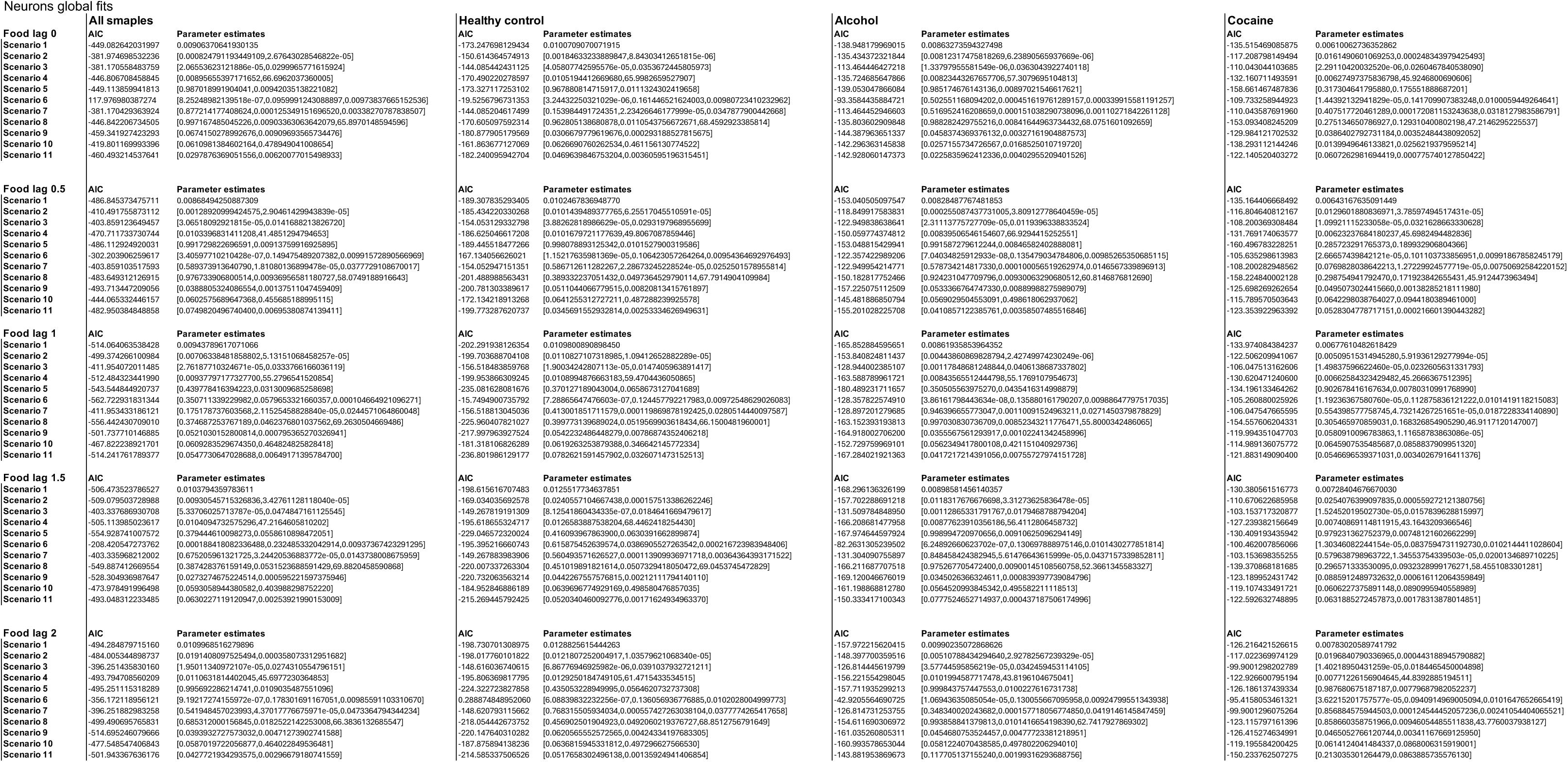

