## Supplementary material for "Effect of Alcohol and Cocaine Abuse on Neuronal and Non-Neuronal Cell Turnover in the Adult Human Hippocampus"

**Mathematical modelling of hippocampal non-neuronal and neuronal cells**

*Global modelling and individual modelling strategies*

The measured $^{14}C$ value is an average concentration from the cells that are present at the time of collection. It is a result of a cumulative incorporation of $^{14}C$ during cell division at different timepoints within the cells still present at collection. Using mathematical models, we can explore different biological scenarios of cell turnover and test whether these scenarios are compatible with our measured data using two strategies: *global modelling* and *individual modelling*.

For *global modelling*, we simulate different cell birth and death processes that affect the average cell age over time and calculate a cell age distribution for each individual for each simulation. This cell age distribution is integrated over the atmospheric $^{14}C$ bomb curve and we calculate what the $^{14}C$ concentration would be with that age distribution. The modelled $^{14}C$ values are then fitted against the measured $^{14}C$ and a goodness of fit is determined based on the cumulative standard error between the modelled and measured $^{14}C$ values. We use Markov Chain Monte Carlo (MCMC) algorithms to simulate thousands of different parameter estimates, to find the best parameters. For *individual modelling*, we have a fixed model with only one parameter to be fitted. We can then estimate the best parameter fit for each individual sample and compare the estimates between the groups, allowing for more variability in this free parameter.

*Equations*

We further developed previously established models where the birth and death rates are represented by partial differential equations (PDE) and where the solution to the equation represents a cell age distribution at any given time in life. In this PDE the density of cells (n, cells/year) depends on the age of the subject (t, in years) and the age of cells (a, in years)[1,2]. There is an initial condition ($f(a)$), representing the initial distribution of cell ages.

$$n(0,a) = f(a), \qquad a > 0$$

Cell birth is:

$$n(t,0) = g(t), \qquad t \geq 0$$

Cell death is represented by $(\gamma)$

$$\frac{\partial n(t,a)}{\partial t}+\frac{\partial n(t,a)}{\partial a}=-\gamma(t,a)n(t,a)$$

The total cell number at time t is

$$\int_0^t n(t,a)da.$$

We then integrate the PDE with the measured atmospheric $^{14}$C levels. If $K(x)$ denotes the atmospheric $^{14}$C level at calendar year $x$, a cell of age $a$ collected at year $y$ would have a corresponding DNA $^{14}$C content $K(y-a)$. Therefore, the measured $^{14}$C concentration in a sample should be the average over all cells, i.e. the integral of the cell density against the atmospheric $^{14}$C.

$$C_{average}=\frac{\int_0^t n(t,a)K(y-a)da}{N(t)}$$

*Description of mathematical models*

We modelled 8 and 11 different biological scenarios, for the non-neuronal population and neuronal population respectively (Supplementary Table 2). The models were structured around two main concepts that affect average cell age: 1. Cell turnover, which is the process of a cell dying and being replaced by a new cell and 2. Cell death without being replaced, which if it is selective towards cells of a specific age will affect the average age. The contribution of these two concepts to the average cell age depend on a few processes which are described below.

Taking hippocampal neurons as an example, different cell processes affect the extent of cell turnover and selective death and thereby the average cell age (Supplementary Fig. 3a). Cell turnover of neurons, adult neurogenesis, only occurs in the dentate gyrus (DG) and not within the cornu ammonis (CA) regions, which are also collected for carbon dating with the neuronal nuclei marker NeuN. The proportion of DG to CA neurons will affect the contribution of adult neurogenesis to the average age as well as the number of cells being exchanged each year. In addition, in rodent models, adult neurogenesis declines with age which could also affect the average age of the cells (Supplementary Fig. 3a).

Less is known about selective loss of cells of a particular age within the human hippocampus. Stereology data suggest that more CA neurons are lost than DG with age, resulting in a selective loss of old neurons with time (Supplementary Fig. 4). In addition, one study suggest that CA neurons are more vulnerable to cell death in individuals with alcohol abuse, however other studies have not been able to demonstrate this selective loss[3,4].

To test models with the different processes affecting cell turnover or selective death described above, we built general models consisting of up to 4 main components and up to 6 different parameters that can be estimated (Supplementary Fig. 3b). Component 1 describes the populations and whether they are subject of turnover or not, a renewing population and a non-renewing population. Parameter 1: *renewing fraction,* is the estimated proportion of cells that are subject of turnover. Component 2 describes the cell turnover dynamics within the *renewing fraction*. The cell turnover dynamics are determined by parameter 2: *turnover rate,* which is the estimated rate of exchanged cells in percent per year within the *renewing fraction* and parameter 3*: turnover decline rate* in the models where turnover changes over time.

For the neuronal population we modeled more complicated models adding a third component that allows for a change in *renewing fraction* over time. In this model, cells from the DG population become non-renewing with time, limiting cell turnover to a smaller and smaller pool. The cells becoming non-renewing could be set to be selective to only prenatal cells or to postnatal cells. Parameter 4: *DG non-renewing rate*, is the rate of which cells within the renewing DG moves to the non-renewing state. The effect of adding this component is that we can allow for cell turnover to decline, not only by declining cell *turnover rate,* but by reducing the *renewing fraction,* the difference between these two strategies is that by reducing the *renewing fraction*, we can select what cells of a certain age are being exchanged and not (Supplementary Fig. 3b).

Components 1-3 and parameters 1-4 make up the backbone of the cell turnover concept with increasing complexity. The selective cell loss concept could be explored as a further complexity as a 4$^{th}$ component. Here we allowed cells within the non-renewing DG population to die or cells within the non-renewing population (CA neurons) to die selectively (Supplementary Fig. 3b) according to stereology data from 5 different studies (Supplementary Fig. 4)[4–8]. We can combine the different core structures and parameters to create different scenarios with increasing complexity (Supplementary Fig. 3b). In total 8 scenarios composed of only components C1 and C2 with increasing complexity, defined as number of parameters and were used for modelling of both non-neuronal and neuronal populations. For the neuronal population, we developed 3 highly complex scenarios composing of all four components and parameters (see Supplementary Table 2).

Each model provides a cell age distribution over time, where the different parameters affect the cell age distribution. To test whether the model is compatible with our data, we take the cell age distribution and integrate this matrix over the atmospheric $^{14}C$ curve from the date of birth to the date of collection, which results in an modelled $^{14}C$ value. We can then test through model fitting the goodness of fit of the modelled values.

*Purity correction of measured $^{14}C$*

The purity of the FACS sorted nuclei was measured. For the non-neuronal population the purity was always >99%. We therefore did not purity correct the non-neuron population. The neuron population was corrected with the following equation:

1) $Corrected\ ^{14}C_{neurons} = (measured\ ^{14}C_{neurons} - measured\ ^{14}C_{non\text{-}neurons}) * (impurity_{neuron}/100)) * 100/purity_{neuron}$

Some subjects did not have an estimated $^{14}C$ value for the non-neuronal population. These were corrected with calculated $^{14}C$ values based on the best fit model for non-neurons (see below). For 4 individuals the purity was not recorded and then an average purity of 96 % was used for purity correction.

*Setting weights, food lag and atmospheric bomb curve*

When modelling the $^{14}C$ data a few technical aspects that might affect the result of the fitted model need to be considered. Firstly, due to the dynamic $^{14}C$ atmospheric curve, small differences in parameter values have dramatically different effects of the estimated $^{14}C$ value depending on when an individual was born. For example, changes in annual turnover creates smaller $^{14}C$ changes in an individual born before 1955 than in an individual born right on the top of the bomb peak as the individual born before 1955 has had longer period of a non-dynamic $^{14}C$ incorporation. Secondly, individuals are also differently sensitive to contemporary contamination depending on when they were born and the mass of the sample. Although we can rule out significant general contemporary carbon contaminations (e.g. from proteins) to our samples using DNA standard controls and blank samples, it is not possible for us to completely rule out any individual contamination to each sample. Samples with small DNA mass theoretically are more affected by small contaminations than samples with larger mass and the AMS measurement becomes less precise, as can be seen by the calculated standard deviation of each measurement. Lastly, multiple studies have shown that there is a lag in $^{14}C$

concentrations in human tissues. Human blood serum samples has been shown to have a deviation of -1.5 ± 0.7 years[9].

Thus, in order to take into account these technical influences we calculated a weight for each individual datapoint based on 1. Sensitivity to contamination 2. Sensitivity to small changes in turnover 3. mass and 4. AMS measuring error. We also explored the different models with different lags of 0, 0.5, 1, 1.5 or 2 years to infer which delay fits best with our data. Our cohort also includes cases from both the northern hemisphere and the southern hemisphere which has slightly different measurements in atmospheric $^{14}C$ values. For our models we assigned for each case a northern or southern hemisphere atmospheric bomb-curve.

*Scenarios and model selection*

We systematically tested different scenarios of turnover and cell death with increasing complexity to find models that fit well with our control data. We first assumed only one population where all cells are subject to turnover and the level of complexity increased with 4 different scenarios of turnover dynamics. The second level of complexity was exploring all the 4 different turnover dynamics scenarios but allowing for a renewing population and a non-renewing population. These 8 models were tested for both neurons and non-neurons. For the neuronal models we added a third level of complexity, allowing for the third DG non-renewing population and cell loss based on stereology data (Supplementary Table 2)[5–8,10].

To explore the best fit turnover scenario, we performed MCMC simulations of all samples in each group. This gives us information on what turnover dynamics fit best with all samples together, using global fitting. The best fit scenario was chosen based on a calculated Akaike information criterion (AIC). The lower AIC value the better the fit. We ranked the fit of the scenarios according to their AIC value. The scenario with the top rank (lowest AIC) was then compared to the scenario with the second best rank. If the top scenario was significantly better ($p < 0.05$) we accepted that scenario as the best fit scenario. If the top scenario was not significantly better than the top 2 scenario we chose the scenario of less complexity and compared this with the top 3 scenario etc. If models parameter estimates resulted in a more complex model to be very similar to a less complex model, e.g. if a turnover decreasing rate was close to 0, we automatically selected the less complex model. If a model had unrealistic estimated values, we did not accept these estimates but instead performed individual turnover fits.

*Setting priors*

Due to the dynamics of atmospheric $^{14}C$, there is for some samples born before the peak, two solutions of *turnover rates* to a $^{14}C$ value (Supplementary Fig. 7). Small changes in low *turnover rates* results in larger differences in $^{14}C$ values that changes in high *turnover rates*. For *global modelling* of the neuronal population we found that this lead to a selection of high turnover solutions, which are not realistic. The *renewing fraction* parameters were more stable and could be used for further modelling of individual *turnover rates*. However when fixing these *renewing fractions* when performing individual fits, many samples could not be fitted as the require slightly larger *renewing fractions*. Therefor we set a prior to all our MCMC models, to put higher likelihood on higher *renewing fractions* and lower *turnover rates* for the neuronal population to estimate a global fit. This did not result in a selection of low turnover solution, but slightly increased the *renewing fraction* so that we could investigate individual *turnover rates* of the samples.

*Investigating turnover dynamics in hippocampal non-neurons*

We calculated weights at food lag 0, 0.5, 1, 1.5 and 2 and found that a food lag at 0.5 had most homogenous weights (Supplementary Table 3). In order not to allow for single samples to drive the models we therefore chose these weights for further modelling. To explore the effect of food lag on improving the general fit we performed Monte Carlo simulation on the models for the control group, ethanol group and cocaine group respectively (Supplementary Table 4). We found, by comparing AIC that a food lag of 0.5 generally improved the fit for most models and therefore continued the analysis with 0.5 food lag.

The model with the best AIC for the all three groups was scenario 5 (Supplementary Table 4). This model has two parameters: *renewal fraction* and *turnover rate* that is constant throughout life (Supplementary Table 2). This scenario was significantly better than the next best model with less complexity, scenario 1, for the healthy control and alcohol groups (log likelihood test, $p< 0.0001$ for healthy control group and $p= 0.0092$ for alcohol group). Scenario 5 was not significantly better than scenario 1 for the cocaine group (log likelihood test, $p> 0.05$). MCMC modelling demonstrated very similar parameter fits for the healthy control, alcohol and cocaine groups (Supplementary Fig. 5), why we decided to do further analysis using scenario 5 for all three groups.

As we could not detect any clear difference between the three groups using *global fit modelling*. We performed *individual fit modelling* on each individual sample. Using the model structure of scenario 5, we fixed the *renewing fraction* and estimated the individual *turnover* *rates*. There was a significant correlation between annual *turnover rates* and age for the healthy control ($F[1, 25]= 22.49$, $R2= 0.5$, $p< 0.0001$, linear regression) and cocaine ($F[1, 14]= 5.78$, $R2= 0.3$, $p= 0.0307$, linear regression) groups, but not the alcohol group ($F[1, 13]= 0.78$, $R2= 0.06$, $p= 0.3939$, linear regression). There was no significant difference between the individual annual *turnover rates* when comparing the three groups (Fig. 2a, $F[5, 52]= 3.990$ $R2= 0.28$, $P= 0.0039$, age was significant: $F[1, 52]= 16.66$, $p= 0.0002$. Group effect was not significant: $F[2, 52]= 0.2688$, $p= 0.7654$, multiple linear regression).

A different way to evaluate a difference in cell turnover, is to compare the difference in individual chronological age and the average cell age, further referred to as *delta age*. The variability in *delta age* will be less as it is limited by the non-renewing population that will remain old, which could then possible make it possible to detect a difference. Based on the individual *turnover rates,* we modelled an average cell age and could calculate the *delta age*. There was a significant correlation between non-neuronal *delta age* and age for the healthy control ($F[1, 25] = 25.00$, $R2=0.5$, $P<0.0001$, linear regression) and cocaine ($F[1, 14]= 13.39$, $R2= 0.49$, $p= 0.0026$, linear regression) groups, but not the alcohol group ($F[1, 13]= 0.98$, $R2= 0.07$, $p=0.34$, linear regression). There was no significant difference between the three groups (Fig. 2b, $F[3, 55]= 201.7$, $R2= 0.92$, $p < 0.0001$. Age was significant: $F[1, 55]= 278.0$, $p< 0.0001$. Group effect was not significant: $F[2, 55]= 0.5329$, $p= 0.5899$, multiple linear regression).

There might be a common cell turnover mechanism of addiction for both alcohol and cocaine, why we combined the two groups into one addiction group. Again there was a significant correlation with age, but no significant difference between the larger addiction group and healthy controls (Supplementary Fig.6, $F[2, 56]= 305.4$, $R2= 0.92$, $p < 0.0001$. Age was significant: $F[1, 56]= 280.8$, $p< 0.0001$. Group effect was not significant: $F[1, 56]= 0.6353$, $p= 0.4288$, multiple linear regression). To determine the amount of reduction in turnover throughout life that would be required for us to be able to detect it, we reduce individual turnover with 30 and 40% and found a significant difference in *delta age* after a reduction of 40% (Supplementary Fig.6, $F[2, 56]= 256.1$, $R2= 0.90$, $p < 0.0001$. Age was significant: $F[1, 56]= 322.4$, $p< 0.0001$. Group effect: $F[1, 56]= 9.817$, $p= 0.0028$, multiple linear regression). Thus, it appears that our non-neuronal population within our healthy controls are highly variable and a fairly large reduction would be required for us to detect a difference.

As we could not detect any difference between the three groups we modelled them all together. Again, scenario 5 was the best model (Supplementary Fig. 5d). For those subjects missing a measured non-neuron $^{14}C$ value, we calculated a $^{14}C$ value based on the parameters from this scenario and used these values for the purity correction of the neuronal $^{14}C$ value.

*Investigating turnover dynamics in hippocampal neurons*

Weights and scenarios were calculated at a lag set to 0, 0.5, 1, 1.5 or 2 years. We found that food lag was most homogenous and the models had best fit at lag of 1 year which was chosen for further analysis (Supplementary Table 5). We tested the same models as for the non-neuronal population and tested three additional models with more complexity. These models had a *DG-non renewing rate* parameter and selective death according to stereology data. We used the same rule of accepting the best scenario as for non-neurons.

For the healthy controls we found that scenario 5 and more complex scenario 11 had equally good fits. However, while scenario 5 was the best fit mode for alcohol and cocaine group, scenario 11 was not a good fit. We found that in all models with two populations, the *turnover rate* estimates were unrealistically high (Supplementary Table 5). This is due to that many samples before the peak have two turnover solutions to the $^{14}C$ value (Supplementary Fig. 7) and changes in low *turnover rates* results in larger differences in $^{14}C$ values than changes in high *turnover rates*, resulting in a higher error when fitting modeled values against the measured values. Scenario 5 was significantly better than the next best model for all three groups and when analyzing all samples together (Supplementary Table 5, Supplementary Fig.8).

Although the turnover estimates from scenario 5 were unrealistic, the *renewing fraction* estimates remained remarkably stable for a wide range of scenarios in the healthy controls, providing a good estimate for further modelling. We therefore performed individual turnover analysis using the *renewing fraction* from the global fit, but allowing for an individual fit of the *turnover rate*. We could not detect any significant correlation of *turnover rates* and age (Healthy control group: $F[1, 22]= 25.00$, $R2= 0.05$, $p= 0.2822$; Alcohol group: $F[1, 19]= 13.39$, $R2= 0.13$, $p= 0.1093$; Cocaine group: $F[1, 15]= 0.98$, $R2=0.20$, $p=0.0686$, linear regression, Supplementary Fig. 9a) nor any significant difference in individual turnover between the three groups (Fig. 3a, Mann-Whitney U test, $p >0.05$). Based on the individual fits, we modelled the average cell age of each sample and calculated the *delta age*. There was no correlation with age (Healthy control group: $F[1, 22]= 0.59$, $R2= 0.03$, $p= 0.4519$; Alcohol group: $F[1, 19]= 1.04$, $R2= 0.05$, $p= 0.3203$; Cocaine group: $F[1, 15]= 0.49$, $R2= 0.03$, $p= 0.4936$, linear regression, see Supplementary Fig. 9b), nor was there any significant difference in *delta age* between the three groups (Fig 3b, p >0.05m Mann-Whitney U test). We combined the alcohol and cocaine group in a common "addiction group" and compared the difference between this larger group and healthy controls. We could not detect any significant difference in neuronal *delta age* between the healthy control group and the addiction group (Fig. 3c, p= 0.1643, Mann-Whitney U test ).

We considered that our study might be underpowered to find biologically relevant differences with our sample size, as the healthy control interindividual variance in individual turnover and *delta age* was very high. We modelled the required decrease in individual turnover that would be required for us to detect a significant difference in *delta age* and found that only a 15% decrease would be required (Fig 3c, Mann-Whitney U test, p= 0.0289). The median individual *turnover rate* of healthy controls and abuse group with 15% reduction was 1.7% and 1% respectively. Thus we would not be able to exclude an individual turnover median difference up to 40% over life within the abuse group.

This data analysis suggest that it is not likely that reduced neurogenesis throughout life is a underlying mechanism of addiction vulnerability, but we are also limited in the ability to detect turnover differences throughout life. The sensitivity is reduced even further when looking at shorter periods of time. Alcohol and cocaine can in animal models reduce neurogenesis after administration. If there would be a significant reduction of as a result of substance intake, the duration of substance intake should correlate with the *delta age*. We did not have any data on the abuse period for the cocaine group, but had data from 18 individuals in the alcohol group, which had a median duration of alcohol abuse of 20 years. We could not detect a significant correlation between alcohol use in years and person *delta age* (Fig. 4a, r= -0.3028, p= 0.2526, Spearman correlation). We modelled the decrease in individual *turnover rates* the last 20 years of life required for us to detect a difference. We found that a 40% decrease of individual *turnover rates* the last 20 years within the alcohol group would be required for us to detect a significant difference in *delta age* (Fig. 4b, 20% reduction: p= 0.0859; 30% reduction: p= 0.0365, Mann-Whitney U test).

It is theoretically possible that selective death of old neurons might mask a reduction in adult neurogenesis. In one stereology study it was observed that a higher fraction of CA neurons compared to DG neurons within younger subjects and that there was a significant difference compared to health controls[3]. However, the difference between controls and the alcohol group was not significant in older age, possibly suggesting that alcohol speeds up an aging process of cell death after alcohol consumption. Within the combined stereology data used for scenario 9-11 we see that around 25% of CA neurons die throughout life[5–8,10]. We modelled the required decrease of individual turnover, throughout life and the last 20 years of life, necessary for us to detect a significant difference in average cell age while allowing for a 25% death of non-renewing cells. At least 40% reduction in individual turnover throughout life (p= 0.0153, Mann-Whitney U test)  and 60% reduction the lats 20 years (p= 0271, Mann-Whitney U test), would be required for us to be able to detect a difference in *delta age* (Supplementary Fig. 10).
